# Predicting DNA origami stability in physiological media by machine learning

**DOI:** 10.1101/2025.07.18.665506

**Authors:** Judith Zubia-Aranburu, Andrea Gardin, Lars Paffen, Matteo Tollemeto, Ane Alberdi, Maite Termenon, Francesca Grisoni, Tania Patiño Padial

**Affiliations:** Department of Biomedical Engineering, Institute for Complex Molecular Systems, Eindhoven University of Technology, Eindhoven, Netherlands; Interdisciplinary Nanoscience Center (iNANO), Aarhus University, Gustav Wieds Vej 14, Aarhus C, 8000, Denmark; Biomedical Engineering Department, Mondragon University, Arrasate-Mondragon, Spain; The Danish National Research Foundation and Villum Foundation’s Center IDUN, Department of Health Technology, Technical University of Denmark, Kgs. Lyngby, Denmark; Centre for Living Technologies, Alliance TU/e, WUR, UU, UMC Utrecht, Princetonlaan 6, 3584 CB, Utrecht, The Netherlands

**Author notes:** These authors contributed equally (Judith Zubia-Aranburu, Andrea Gardin).

**Keywords:** DNA origami, stability, dynamic light scattering, diffusion coefficient, machine learning

## Abstract

DNA origami nanostructures offer substantial potential as programmable, biocompatible platforms for drug delivery and diagnostics. However, their structural stability under physiological conditions remains a major barrier to practical applications. Stability assessment of DNA origami nanostructures has traditionally relied on image-based and empirical approaches, which are time-consuming and difficult to generalize across conditions. To address these limitations, we developed a machine learning approach for DNA origami stability prediction, based on measurable physicochemical parameters. Using dynamic light scattering (DLS) to quantify diffusion coefficients as a proxy for structural integrity, we characterized over 1400 DNA origami samples under varying physiologically relevant variables: temperature, incubation time, MgCl_2_ concentration, pH, and DNase I concentrations. The predictive performance of the model was confirmed using an independent set of samples under previously untested conditions. This data-driven approach offers a scalable and generalizable framework to guide the design of robust DNA nanostructures for biomedical applications.

## 1. Introduction

Over the past decade, DNA origami nanostructures^1–3^ have emerged as highly promising tools for drugs ^4^ and gene delivery,^5–7^ cell receptor targeting, ^8–11^ or molecular diagnostics.^12–14^ Their unique properties, including high programmability, nanoscale precision, biocompatibility and the capacity for selective targeting of cells and tissues, have enabled the fabrication of increasingly sophisticated nanostructures, expanding the frontiers of DNA nanotechnology.^15^ However, a key challenge remains in traslating these systems into biological and/or clinical context: maintaining structural stability under physiological conditions.^16^

Stability, defined as the ability of the structure of a DNA origami to persist under environmental stressors, has been extensively studied.^17–19^ Environmental factors such as temperature, ionic strength, pH variations and enzymatic degradation by nucleases are known to compromise the integrity of these nanostructures.^20^ Multiple strategies have been developed to enhance stability, *e.g.*, covalent chemical modifications,^21,22^ protective surface coatings,^23^ and fine-tuning of assembly parameters. For instance, covalent cross-linking^24^ and the use of stabilizing agents, like polyethylene glycol (PEG),^25,26^ can improve the structural integrity of DNA origami in harsh environments. Furthermore, encapsulation of DNA origami in protective shells,^27^ such as liposomes,^28^ polymers^29^ or silica coatings,^30^ has also demonstrated protection against de-stabilizing factors.

Nevertheless, most current approaches to improving DNA origami heavily rely on experimental trial and error. These methods are often labour-intensive, require specialized instrumentation, and do not easily scale to diverse physiological conditions relevant for biomedical applications. There is, therefore, a pressing need for a more generalizable, predictive framework to assess DNA origami stability more efficiently and systematically.

In recent years, machine learning (ML), a subset of artificial intelligence that enables algorithms to learn relevant information directly from data, has become an increasingly valuable tool in the molecular sciences, offering the ability to capture complex biophysical relationships in molecular systems from large experimental datasets.^31^ Within DNA nanotechnology, such methods have been applied to optimize nanorobot design, ^32^ predict DNA hybridization kinetics,^33^ improve DNA origami structure identification and characterization,^34^ and enable particle detection and pose estimation.^35^ Similar strategies have also been employed to model thermodynamic stability and folding probabilities in nucleic acid structures such as G-quadruplexes.^36^ Together, these developments demonstrate how machine learning can aid experimental planning and speed-up the design of functional molecular nanostructures.

Motivated by this recent progress, here we develop a machine learning framework to predict the stability of DNA origami structures across different physiological conditions. Our starting point was high-quality, quantitative data, generated by dynamic light scattering (DLS). DLS offers several advantages: it is high-throughput, quantitative, broadly accessible, and less resource-intensive than traditional stability assays. Techniques such as gel electrophoresis^37^ and circular dichroism^38^ are simple but qualitative, while others like atomic force microscopy,^39^ transmission electron microscopy^40^ and fluorescence resonance energy transfer assays^41^ can provide quantitative data but are low-throughput and require specialized equipment and expertise. Using DLS, we systematically tested the stability of DNA origami nanostructures at different physiological conditions, including temperature, incubation time, MgCl_2_ concentration, pH and DNase I concentration (Figure 1). We compiled a dataset of over 1400 measurements across a range of geometries, which we used to develop a ML model capturing the relationship between environmental conditions, nanostructure shape and stability (Figure 1).

**Figure 1.**
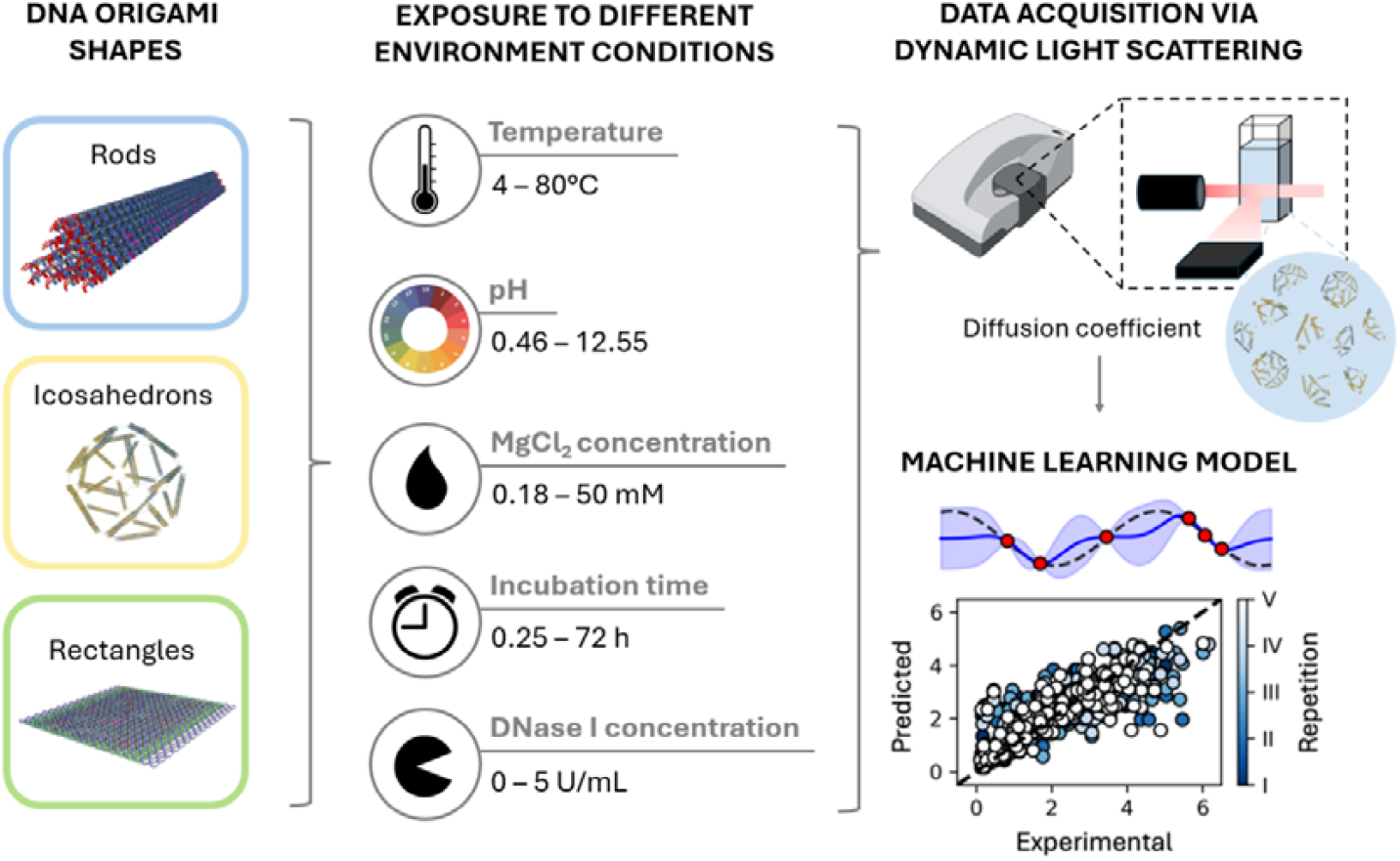
Study workflow. Starting, from left to right, the assembly of DNA origami nanostructures, in three different shapes (rods, icosahedrons and rectangles); their exposure to temperature, pH, MgCl_2_ concentration, incubation time and DNase I concentration values; analysis of their stability by dynamic light scattering and generation of the ML model.

The predictive accuracy and generalizability of the model was confirmed via prospective validation. Our results underscore the developed machine learning model as a practical tool for rational design, ultimately enabling the development of DNA origami structures with improved stability profiles, to be tailored to specific biomedical contexts and applications.

## 2. Results and discussion

### 2.1 DNA origami assembly and characterization

We assembled three DNA origami nanostructures, spanning various sizes and shapes, as previously reported in literature, including: (a) icosahedrons,^42^ with a diameter of 45 nm, (b) rectangles ^43^ of size 75 × 100 × 2 nm, and (c) rods,^44^ with a diameter of 15 nm and length of 150 nm. Once assembled, the DNA origami nanostructures were purified to eliminate unbound ssDNA origami staples. To confirm correct purification, a gel electrophoresis was conducted (Figure 2D). The correct nanostructured assembly was confirmed by AFM (Figure 2B), DLS (Figure 2C) and gel electrophoresis (Figure 2D). In AFM measurements, both rectangles and rods showed a perfectly matching size between experimental and theoretical values. In the case of icosahedrons, a mean lateral diameter of 80 nm and maximum height of 6.4 nm was observed, differing from the theoretical value of 45 nm. This could be explained by a “flattening” effect due to the pressure exerted by the AFM tip during imaging procedures. This hypothesis was verified by DLS, which confirmed the correct assembly of the three structures, as all of them displayed an expected hydrodynamic diameter, matching the theoretical value.

**Figure 2.**
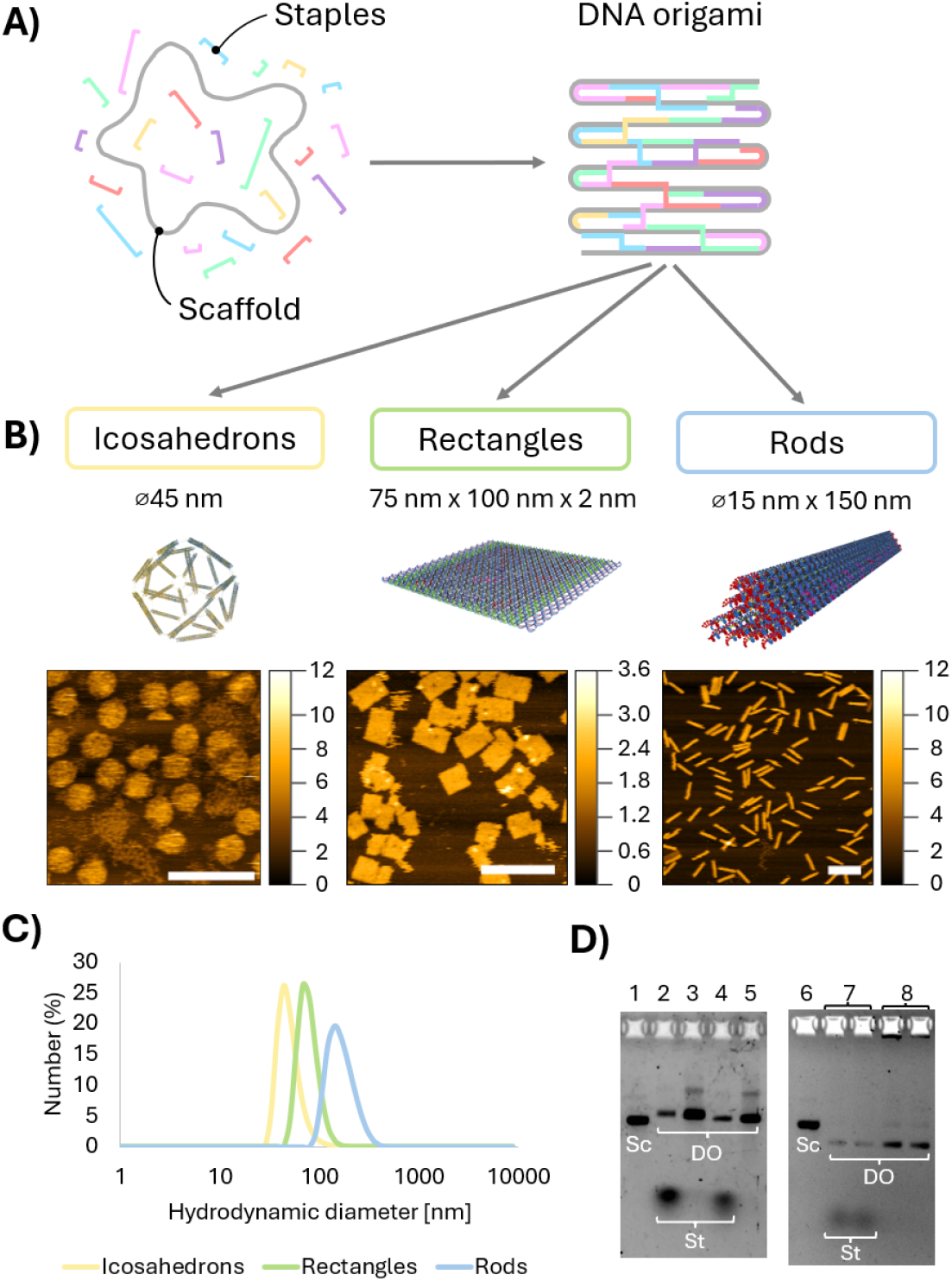
DNA assembly and characterization. (**A**) Schematic representation of the components and formation of DNA origami nanostructures. (**B**) 3D rendered representation of the icosahedrons, rectangles and rods, and their corresponding AFM images (scale bar = 200 nm; heigh values in nm). (**C**) DLS characterization of the three origami shapes. D) GE of purified and unpurified origami shapes, where 1 is M13 scaffold, 2 unpurified icosahedrons, 3 purified icosahedrons, 4 unpurified rectangles, 5 purified rectangles, 6 p7560 scaffold, 7 unpurified rods and 8 purified rods. Sc refers to the scaffold, St to staples and DO to DNA origami.

### 2.2 Generation of DNA origami stability dataset

After DNA origami characterization, we established a quantitative parameter as a proxy for structural stability. We selected dynamic light scattering (DLS) as a main technique due to its rapid, non-invasive and reliable assessment of the DNA origami structural stability, where a high number of DNA origami nanostructures are simultaneously measured.^45^ DLS measures the diffusion coefficient of nano-sized particles, through the decay rate of the intensity autocorrelation function, which reflects the temporal fluctuations in scattered light caused by Brownian motion. The Stokes-Einstein equation is then used to calculate the hydrodynamic size of the particles.

In this context, diffusion coefficient and hydrodynamic size both provide valuable, complementary insights into structural integrity and aggregation behaviour of DNA nanostructures. Under stable conditions, these structures exhibit characteristic diffusion coefficient values. Structural degradation and/or disassembly into smaller fragments result in increased diffusion coefficients, while aggregation leads to larger particle sizes, and correspondingly lower diffusion coefficients (Figure 3A). Although hydrodynamic size offers an intuitive metric for comparing expected versus observed structural dimensions, it can be misleading when applied to DNA origami, due to deviations from the spherical geometry assumed by the Stokes-Einstein model. In contrast, the diffusion coefficient is a more fundamental physical parameter, as it directly reflects molecular motion and is less influenced by shape assumptions. Moreover, it is directly measured by DLS and is highly sensitive to subtle structural changes. For these reasons, we adopted the diffusion coefficient as the principal proxy for assessing DNA origami stability.

**Figure 3.**
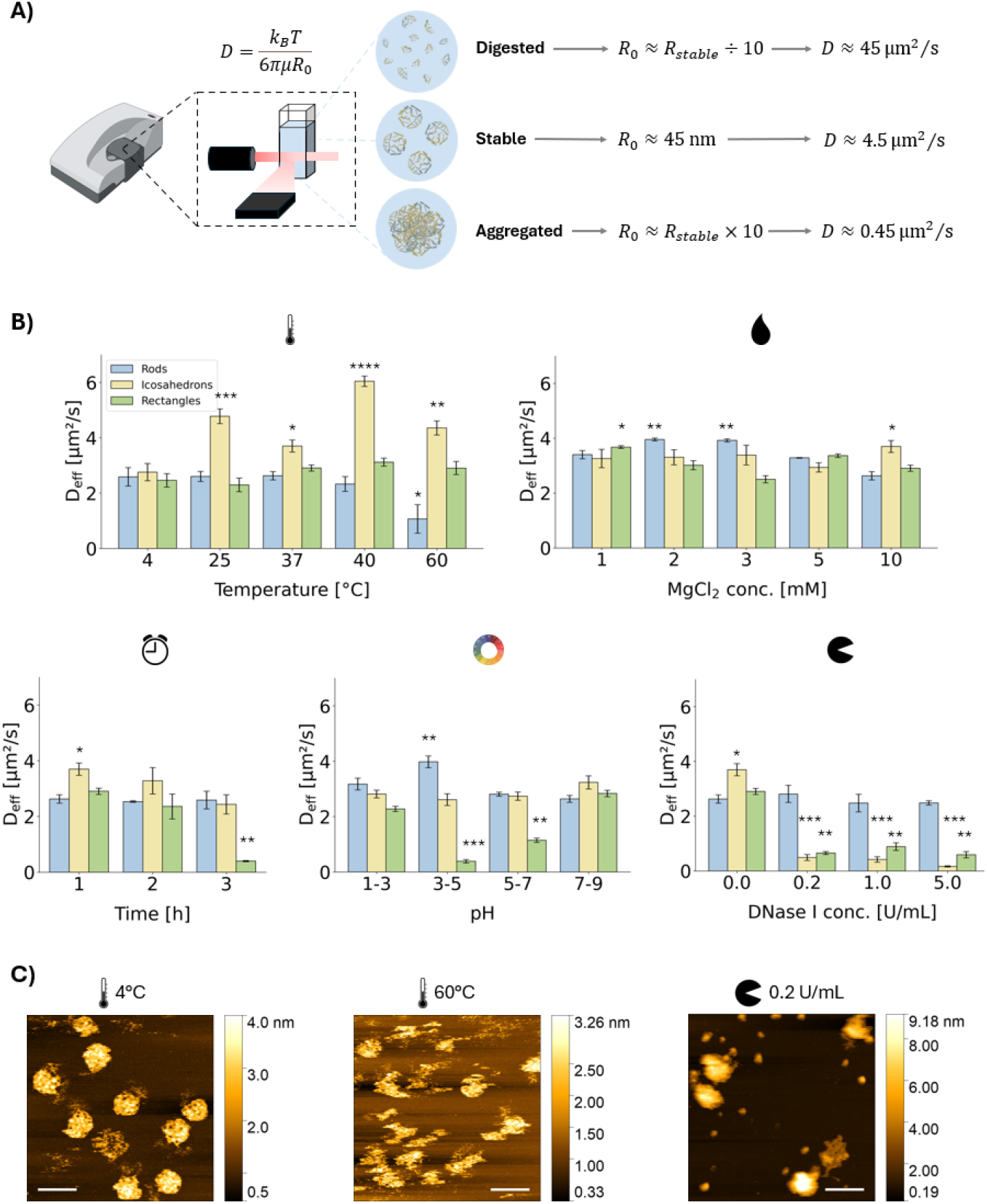
Generation of DNA stability dataset. (**A**) Schematic representation of the change of diffusion coefficient, as measured by DLS, upon destabilization of the DNA nanostructure. The relationship between the radius and diffusion coefficient is provided to aid understanding. *D* is diffusion coefficient, *k*_*B*_ Boltzmann’s constant, *T* temperature and μ solvent viscosity. (**B**) Diffusion coefficient values measured via DLS for each shape at different experimental conditions (temperature, MgCl_2_ concentration, incubation time, pH and DNase I concentration). Error bars represent the standard error of the mean. All p-values were computed from the unpaired t-test with unequal variance with respect to the diffusion coefficient at 4°C; *p-value ≤ 0.05; **p-value ≤ 0.01, ***p-value ≤ 0.001, ****p-value ≤ 0.0001. (**C**) Representative AFM height images corresponding to the control (4°C) and two unstable conditions (60°C and 0.2 U/mL of DNase I) depict the different ways in which loss of structural stability occurs, leading to different diffusion coefficient values (scale bar = 100 nm).

The three distinct DNA origami shapes (i.e., rods, icosahedrons and rectangles) were exposed to key stressors, including variations in temperature, incubation time, MgCl_2_ concentration, pH and DNase I activity. Diffusion coefficients were then measured using DLS to quantify the structural response under each condition (Figure 3B). As a control, we used samples maintained under standard conditions (4°C, 10 mM MgCl_2_, 1 h incubation, neutral pH and without DNase I), which are considered optimal for DNA origami assembly and storage.

We observed a clear shape-dependent effect on structural stability. Rod-shaped nanostructures exhibited the highest stability, with minimal fluctuations in diffusion coefficients across tested conditions. This resilience is likely attributable to their compact design, as illustrated in Figure 2B. In contrast, rectangular structures showed greater susceptibility to instability, particularly under varying pH, incubation time and DNase I activity, whereas temperature and MgCl_2_ concentration had a less pronounced effect. The sharper sensitivity to pH may be explained in terms of protonation or deprotonation of nucleobases under acidic or alkaline conditions, which can disrupt base pairing.^46^ Additionally, the relatively exposed surface area of rectangular DNA origami compared to the other shapes likely facilitates nuclease access, accelerating degradation and leading to complete desintegration.^18^

Among the three, icosahedrons were the most structurally sensitive, which may stem from their less compact design compared to rods and rectangles. Their stability was particularly compromised by elevated temperatures and DNase I exposure. At 4°C, the control samples maintained well-formed icosahedrons (see Figure 3C). However, exposure to higher temperatures induced fragmentation into smaller pieces (Figure 3C), as reflected by increased diffusion coefficients, which could be attributed to a detachment of staple strands from the scaffold. Conversely, DNase I exposure led to the formation of aggregates, as seen in the AFM images (Figure 3C), where bulkier structures replaced the well-defined morphologies observed in the control sample. This aggregation caused a sharp decrease in the diffusion coefficient, approaching zero.

Recognizing that changes in diffusion coefficient can also arise from factors unrelated to degradation (i.e. nonspecific interactions or minor conformational shifts), we complemented DLS measurements with gel electrophoresis (Supporting Figures S1-S3) and some more atomic force microscopy (AFM) images (Supporting Figure S4) to support and confirm our structural findings.

### 2.3 Predicting the diffusion coefficient with machine learning

The final dataset contained 1417 DNA origami nanostructures, encompassing a range of shapes and experimental conditions (Figure 3). This dataset was used to train and validate a machine learning model to predict the corresponding diffusion coefficient. Of the total collected samples, 1317 were allocated for model optimization, training, and testing, while 100 were reserved as a held-out set for final validation, selected using a farthest point sampling strategy to maximize their diversity (Figure 4A). The former set (1317 samples) was further split into training and test subsets with a 70%:30% ratio, repeated across five randomized repetitions (Figure 4B, and Materials and Methods).

**Figure 4.**
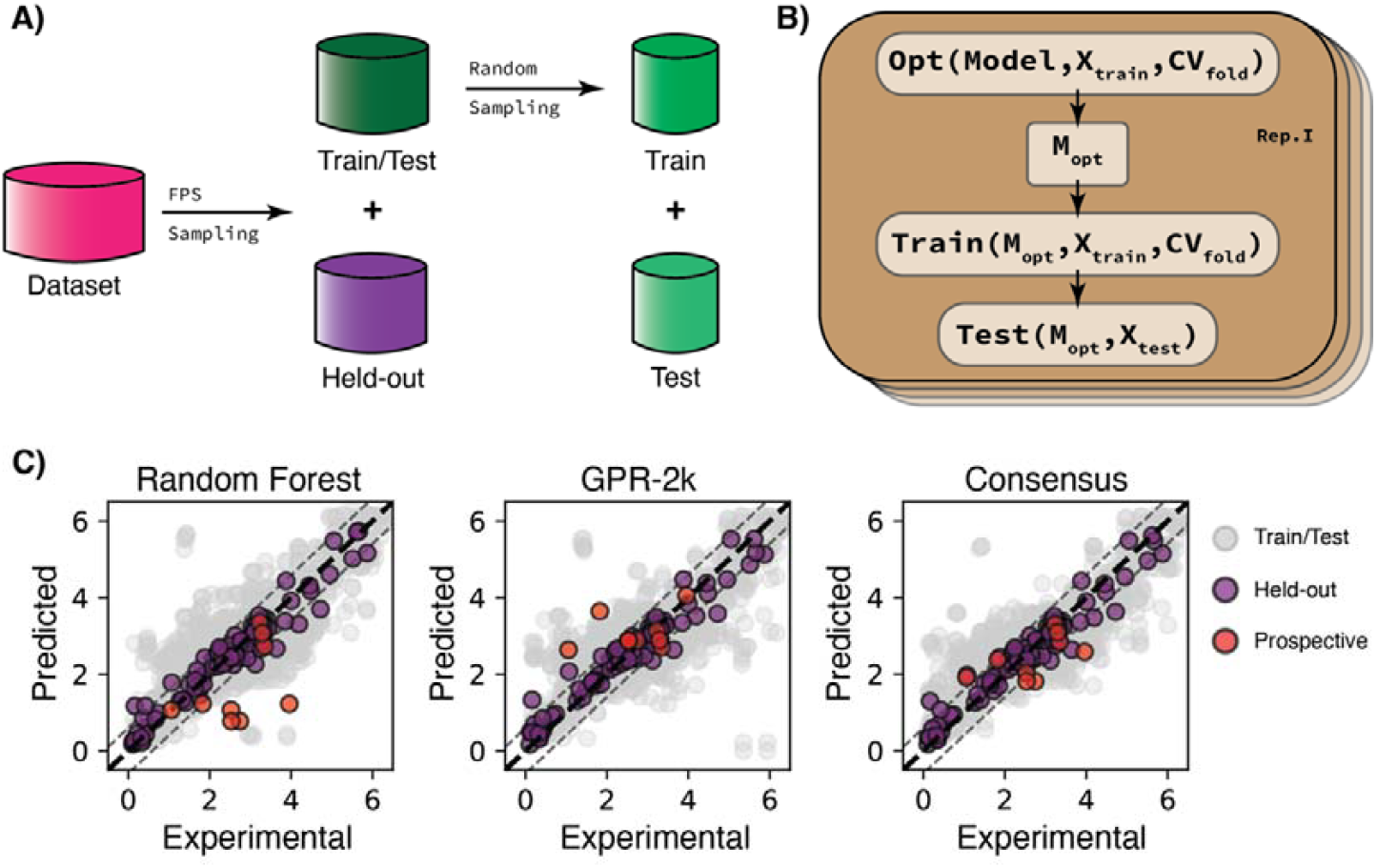
Machine Learning experiments. (**A**) Dataset splitting strategy. A held-out set was sampled using farthest point sampling (FPS), and the remaining data were further split into a training set (70%) and a test set (30%). (**B**) Schematic of the model training and evaluation pipeline. Each repetition of the process begins with an optimization step (*Opt*) where different versions of the model are tested on the training data (*X*_*train*_) using cross-validation (CV_fold_, a technique that splits the data into parts to ensure fair testing). The best performing model (M_opt_) is then trained on the full training data and evaluated on unseen test data (X_test_) to assess its performance. The entire sequence is repeated to ensure reliability of the results. (**C**) Experimental vs predicted diffusion coefficient for Random Forest (RF), Gaussian Process Regressor with 2 kernel functions (GPR-2k), and the consensus model built from merging RF and GPR-2k predictive power. Colors represent the considered dataset (grey points were obtained as a superimposition of five training repetitions). The grey dotted lines delimit an interval of confidence that matches the average variability in experimental measurements.

In this work, we evaluated four well-established machine learning models: (a) Random Forest regressor (RF),^47^ which builds multiple decision trees and uses their consensus to provide a prediction; (b) Extreme Gradient Boosting (XGBoost),^48^ which sequentially builds decision trees in a step-wise process, where every step the prediction is aimed to correct previous errors;(c) Support Vector Regressor (SVR),^49,50^ which identifies hyperplanes that best describe the target response variable; and (d) Gaussian Process Regressor (GPR),^51^ a probabilistic approach that provides predictions alongside an estimation of uncertainty. GPR models rely on kernel functions, which define how input features are combined to make predictions. In this work, we implemented two variations: GPR-1k, which employs a single kernel function, and GPR-2k, which linearly combines two kernel functions, potentially capturing more complex relationships (Eq. 2-3).

Model performance was assessed using the Root Mean Squared Error (RMSE, Eq. 4), which quantifies the error in the predicted diffusion coefficient (expressed in µm^2^/s). Across the repeated training-test splits (Figure 4C and Table 1), RF, XGBoost, and GPR-2k showed the best performance (RMSE values ranging from RMSE = 0.68 µm^2^/s to RMSE = 0.81 µm^2^/s). Such values were in the same range of magnitude than the experimental error across repeated measurements (estimated to be equal to ±0.42 µm^2^/s in our experimental setup) and hence considered as satisfactory for a predictive model. Notably, the combination of kernel functions in GPR-2k led to an improvement in performance compared to using a single-kernel, possibly due to the ability to better capture both local and global patterns. SVR exhibited suboptimal performance. As a result, only RF, XGBoost, and GPR-2k were selected for the next evaluation phase, using the held-out data (100 samples). Additionally, we created a *consensus* model that combines the predictions of RF and GPR-2k by averaging their outputs, aiming to leverage their complementary strengths and improve robustness. On the training and test sets, the consensus model achieved respectively comparable and lower RMSE scores than the individual models (Table 1).

**Table 1.** Model performance. Root Mean Squared Error (RMSE [µm^2^/s]; the lower, the better) obtained on each set split. For training and test set, the average RMSE, and its standard error are reported over five splitting repetitions. Best and second-best performance for each set is reported in boldface and with underlying, respectively. A consensus model was obtained by averaging the predictions of RF and GPR-2k.

| Set | RF | SVR | GPR-1k | GPR-2k | XGBoost | Consensus |
| --- | --- | --- | --- | --- | --- | --- |
| Training | **<u>0.68±0.07</u>** | 1.27±0.12 | 0.89±0.05 | 0.81±0.07 | <u>0.74±0.04</u> | 0.77±0.03 |
| Test | 0.72±0.04 | 1.25±0.20 | 0.93±0.03 | 0.75±0.05 | <u>0.67±0.04</u> | **<u>0.66±0.04</u>** |
| Held out | **<u>0.34</u>** | *n.a.* | *n.a.* | 0.36 | <u>0.35</u> | <u>0.35</u> |
| Prospective | 1.39 | *n.a.* | *n.a.* | <u>0.86</u> | 1.44 | **<u>0.73</u>** |

The held-out set was used to evaluate model performance on datapoints that fell within the range of the tested experimental conditions, but that were not used to train the model. All four models performed similarly on this set, yielding RMSE values around 0.35 µm^2^/s (with negligible differences when considering the experimental error). Interestingly, the RMSE values on the held-out set were lower than those from the repeated train-test splits (Table 1). This may arise from the farthest point sampling strategy, which ensured a representative selection of data points, as well as differences in shape proportions within the held-out set. Additionally, the smaller sample size could have contributed to reduced variability in RMSE estimates. Taken together, these factors suggest that while the held-out set provides a useful benchmark for model performance, the results from the repeated test splits have to be taken into consideration and remain crucial for assessing overall generalization ability.

### 2.4 Prospective model validation in new experimental conditions

To further evaluate the performance of the top-performing models under previously untested conditions, we designed nine new experiments positioned at the boundaries or beyond the ranges of the original dataset. These results revealed a clear advantage for the GPR-2k and the consensus model, which outperformed both Random Forest and XGBoost in terms of predictive accuracy (Figure 4C; Table 1; Supplementary Figures S5–S7). Among the models tested, GPR-2k was the only one to maintain consistent RMSE values across the training/test, held-out, and prospective sets. The consensus model, constructed by averaging predictions from GPR-2k and Random Forest, achieved the lowest RMSE on the prospective data, indicating that combining probabilistic and tree-based approaches can enhance robustness and generalization, especially under extrapolative conditions. Notably, only 3 out of 9 predictions (Figure 5, samples 1, 5, and 9) scored values outside the experimental variance, further supporting the reliability of the combined modelling strategy.

**Figure 5.**
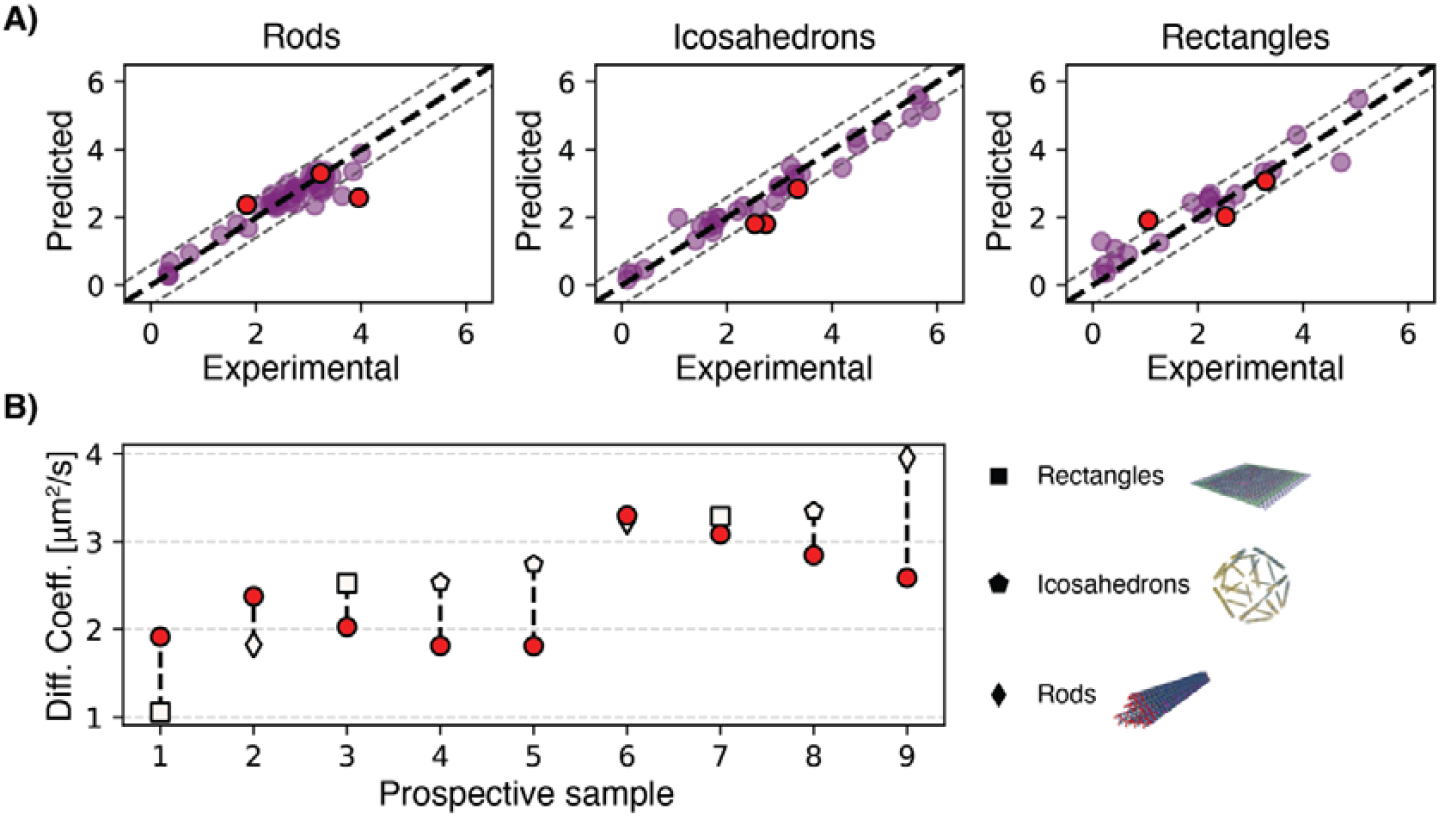
Prospective application of the consensus model. (**A**) Experimental vs predicted diffusion coefficients, divided by shape (left to right). In each correlation plot, see-through points in the background represent the held-out set (as shown in Figure 4), and opaque points in the foreground are the prospective dataset. The grey dotted lines delimit an interval of confidence that matches the average variability in experimental measurements. (**B**) The tested prospective samples ranked based on the extent of their recorded diffusion coefficient (increasing from left to right). The dotted vertical line represents the distance between the predicted diffusion coefficient (circles) and the real experimentally recorded value (shaped data points).

The surprising difference in performance of the consensus model could be attributed to advantageous characteristics native of GPR. Unlike the other two tree-based methods, GPR produces smooth predictions, avoiding abrupt changes when changing experimental conditions, hence being potentially more aligned with the physical nature of the problem.^52,53^ In contrast, RF and XGBoost excel in performance when tabular data is present (e.g., the one-hot encoded origami shapes). Finally, GPR provides probabilistic predictions, offering both a mean value and an uncertainty estimate, which enhances interpretability and allows for more informed decision-making prospectively.^51^ Taken together, these results and considerations highlight the consensus model as a good strategy to explore the stability of DNA origamis prospectively.

### 2.5 DNA origami stability threshold determination

Once we established an efficient method to predict the diffusion coefficient of DNA origami particles, we shifted our attention back to investigating their stability using the coefficient values as a key discriminant. To the best of our knowledge, this represents the first attempt to define the stability of these supramolecular structures based on diffusion coefficient ranges.

To qualitatively categorise the predicted diffusion coefficient values with stable or unstable states, we established thresholds values for each particle shape. Based on our previous experimental findings, reported in Figure 3, and the observed shape-wise coefficient ranges, we defined both lower and upper thresholds for the stability of the nanostructures. Specifically, the first and third quartile values of the 4°C sample of each shape were established as lower and upper threshold values, respectively (see Table 2). These thresholds enabled a qualitative framework to determine the nanostructure stability by only comparing the predicted diffusion coefficient values to shape-specific stability ranges.

**Table 2.** Determined lower and upper threshold values of the diffusion coefficient corresponding to stable rods, icosahedrons and rectangles, based on our experimental findings.

| Shape | Lower threshold | Upper threshold |
| --- | --- | --- |
| **Rod** | 1.9 $\mu\text{m}^2/\text{s}$ | 3.3 $\mu\text{m}^2/\text{s}$ |
| **Icosahedron** | 1.8 $\mu\text{m}^2/\text{s}$ | 4.0 $\mu\text{m}^2/\text{s}$ |
| **Rectangle** | 1.4 $\mu\text{m}^2/\text{s}$ | 2.8 $\mu\text{m}^2/\text{s}$ |

While these thresholds remain qualitative in nature and specific to our experimental set of screened variables, they mark an important step in advancing the understanding of DNA origami particle stability. Out of the nine new experimental conditions tested only two resulted in a significantly different diffusion coefficient values, or stability measurement, demonstrating the potential of a machine learning model in providing new insight into the stability of such nanostructures.

## 3. Conclusions

In this study, we demonstrated the feasibility of using the diffusion coefficient as a quantitative proxy for assessing the structural stability of DNA origami nanostructures. Through the combined use of gel electrophoresis, dynamic light scattering and atomic force microscopy, we established a strong correlation between changes in diffusion coefficient and structural transitions associated with assembly and disassembly processes. Our analysis encompassed three distinct DNA origami geometries (rods, icosahedrons and rectangles), supporting the generalizability of this approach across a broad range of shapes.

We systematically evaluated the stability of these structures under a variety of environmental conditions, including temperature, MgCl_2_ concentration, incubation time, pH, and DNase I concentration. DLS measurements revealed shape-specific responses to these stressors. Rods exhibited the highest overall stability, while all structures showed pronounced destabilization under extreme conditions, such as temperatures above 40°C, prolonged incubation (>3h), pH outside the range of 6-8, or DNase I concentrations exceeding 0.2 U/mL.

The high-throughput capacity of DLS enabled the collection of more than 1400 data points, forming a robust foundation for training a predictive machine learning model. We evaluated four regression algorithms, each with distinct advantages. Among them, a consensus model built from Gaussian Process Regressor (GPR) and Random Forest (RF) outperformed others, owing to its joint flexibility, smooth interpolation, and strong predictive performance even with relatively limited training data. Notably, the consensus model was validated on an independent prospective dataset consisting of conditions outside the training distribution, achieving excellent agreement with experimental results.

To further enhance the interpretability of the predictive model’s output, we introduced shape-specific stability thresholds, defined by the lower and upper quartiles of the diffusion coefficient distributions. While these thresholds are inherently tied to our experimental setup and should be interpreted qualitatively, they represent an innovative step toward a more structured, quantitative framework for assessing supramolecular stability. Furthermore, the machine learning model’s utility extends beyond this study, as it can be used iteratively to refine these thresholds or extend to new structural contexts, offering a versatile tool for future investigations.

Altogether, this work lays the foundation for data-driven strategies in evaluating and optimizing DNA origami nanostructures. Such approaches hold promise for deepening our understanding of structural behaviour under physiological conditions and accelerating the design of more robust DNA-based materials for biomedical applications.

## 4. Materials and methods

### DNA origami formulation

#### DNA origami assembly and purification

The DNA origami nanostructures (rods, icosahedrons, rectangles) were assembled by mixing together the scaffold (p7560 for rods; M13mp18 for spheres and rectangles) (Tilibit) and staple strands (Integrated DNA Technologies) in one-pot reaction mixture, in the presence of monovalent and divalent cations, followed by thermal annealing the reaction mixture in the T100™ Thermal Cycler (Bio-Rad). Staple strand sequences are given in Table S1, Table S2 and Table S3 for rods, icosahedrons and rectangles, respectively.

After assembly, DNA origami rods underwent purification via polyethylene glycol (PEG) precipitation by mixing the precipitation buffer (15% PEG8000 (w/v), 5 mM Tris-HCl, 1 mM EDTA, 1,010 mM NaCl (pH 8.0)) in a 1:1 ratio with the DNA origami reaction mixture ^54^. Spheres and rectangles were purified by ultracentrifugation via 100 kDa MWCO 0.5 mL Amicon® Ultra centrifugal filters (Merck Millipore) only for AFM imaging (not for the destabilization assays) due to the inefficiency of PEG precipitation and high rate of origami loss with ultracentrifugation.

The final concentration of the DNA origami nanostructures was determined via the NanoDrop UV-Vis spectrophotometer (Thermo Fisher Scientific) by measuring the absorption at a wavelength of 260 nm. The following molar extinction coefficient values were used: L_rod_: 9.45 × 10^7^ M^-1^cm^-1^; □_icosahedron_: 9.45 × 10^7^ M^-1^cm^-1^; □_rectangle_: 1.24 × 10^8^ M^-1^cm^-1^. All samples were stored at 4°C for a maximum of one week before carrying out the experiments.

#### Sample preparation and destabilization assays

All DNA origami samples to be tested in the destabilization assays were prepared in 1.5 mL Eppendorf tubes to achieve a concentration of 10 nM within a total volume of 100 μL. This was done by using a storage buffer with the MgCl_2_ and DNase I concentrations of interest, followed by the addition of either HCl or NaOH to adjust the pH. The samples were incubated at a predetermined temperature and no shaking for a specific duration between 15 min and 3 days.

### DNA origami characterization

#### DLS measurements

DLS measurements were carried out at the Zetasizer Nano ZSP (Malvern Panalytical) at 25°C and with a backscatter angle of 173°, using a sample volume of 75 µL. After incubation, DNA origami samples were allowed to equilibrate at room temperature for at least 15 min, preventing potential biases in data due to changes in Brownian motion induced by temperature variations. All DLS measurements adhered to a standard operating procedure tailored for proteins, characterised by a refractive index of 1.45 and an absorption value of 0.001, within an aqueous medium with a refractive index of 1.33. Each sample underwent three consecutive measurements, ensuring robust data acquisition and statistical significance.

#### Gel electrophoresis

Agarose gel electrophoresis was conducted using 1.5% agarose gel casted in gel buffer (0.5× TBE, 10 mM MgCl_2_, pH 8.0) and supplemented with SYBR™ Safe DNA Gel Stain. DNA origami samples (10 nM in 10 μL) were loaded with 1.5% (w/v) Ficoll®-400 and let run in gel buffer for 90 min at 75 V in an ice bath. Images were acquired via ImageQuant 800 (Amersham™) and analysed with ImageJ.

#### Atomic Force Microscopy imaging

AFM images of the DNA origami nanostructures were acquired in tapping mode under liquid conditions by means of the Cypher ES Environmental AFM (Asylum Research) using a silicon nitride cantilever BL-AC40TS-C2 (Olympus) with a nominal spring constant of 0.09 N/m. Substrates were prepared by attaching a 1 cm^2^ mica disc (Ted Pella) onto a magnetic puck through double sided tape. Subsequently, mica was cleaved to expose a negatively charged, hydrophilic and atomically flat surface. DNA origami samples were diluted to 2 nM in imaging buffer (1x TAE, 1 mM EDTA, 10 mM MgCl_2_, pH 8.0) prior to their visualization. A 5 μL aliquot of the sample was added onto the substrate, let settle for 30 seconds, followed by the addition of 50 μL imaging buffer. To avoid the formation of bubbles, 50 μL of imaging buffer were also added to the cantilever. All images were analysed using Gwyddion 2.65 software.

### Machine learning

#### Data preparation and splitting

The dataset consisting of 1417 samples and their diffusion coefficients was split into (1) a model development dataset (optimization, training and testing, 1317 samples), and (2) a held-out set (100 samples), via farthest point sampling algorithm. The held-out set did not participate in model development and selection but was used only for the final model validation. The model development dataset was then further split into a training set (70%) and a test set (30%) via random partitioning, for five independent and randomized splitting procedure. The training set was used for hyperparameter selection (via five-fold cross-validation), and the chosen model was validated on the respective test-set. Before testing the final models on the held-out set, the models were re-trained on the full Model Development dataset (1317 samples).

#### Data description and scaling

Each sample was described by: (a) five numerical variables, i.e., temperature of the solution, concentration of MgCl_2_, final pH of the solution, incubation time of the solution, concentration of the DNase I enzyme, and (b) categorical information on DNA origami shape (rectangles, rods, and icosahedrons), which was one-hot encoded into three sparse vectors (having a value of ‘1’ in the column capturing the corresponding shape, and a ‘0’ otherwise). All variables were ‘min-max’ scaled (to obtain values between 0 and 1) using the parameters of the dataset used for model training. For each of the five independent random splitting the scaler was trained only on the training set (70%) and subsequently used to scale the test set (30%) accordingly.

#### Machine learning approaches

- *Gaussian process regression.* Both GPR models were based on a kernel function of this form:

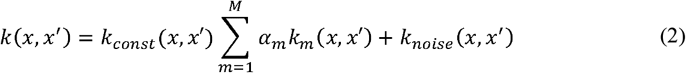

where *k*_const_ represents a constant value kernel function, *k*_m_ represents the kernel function (e.g., RBF, Linear, etc.), *α*_m_ is a constant parameter added for regularization of the kernel function, and *k*_noise_ is the white noise kernel. The function is defined as:

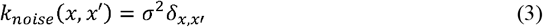

σ^2^is the variance of the noise and 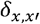 is a Kronecker delta, which equals 1 if, and *x = x^′^*and zero otherwise. In all the kernel functions, the two variables,*x,x^′^*, represent two input feature vectors: is a vector representing one single observation in the dataset,*x*^*′*^ another vector used to compare against in the kernel function. The model relies on these types of functions to define a similarity measure between points, allowing the model to infer values smoothly by con sideri ng how similar a new input is to previously observed data points *x*^*′*^ In Eq. 2,*k*_m_(*x,x^′^*) can take different functional forms: from simple linear, to more complex non-linear ones, and it is crucial to effectively learn from the data (*e.g.*, make predictions). Conversely, the function in Eq. 3, serves as a regularization parameter which adds white noise to the prediction, mimicking the noise of real experiments, avoiding possible overfitting scenarios. In our implementation, we choose two values of *M*: *M* = 1, leading to the usage of one kernel only (GPR-1k), and *kM*(*x,x*^′^) = 2, yielding two linearly combined kernels (GPR-2k). In all cases, _m_ (Eq. 2) was chosen as a radial basis function.
- *Other machine learning algorithms.* Random forest regressor, support vector regressor and XGBoost were applied via standard implementations (*see* Software and Code).
- *Consensus model.* The consensus model was built by merging the RF and GPR-2k models together. The outcome prediction is computed as a simple average of the two predictions.

#### Performance quantification

Model performance was quantified via the Root Mean Squared Error (RMSE), computed as follows:

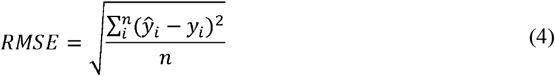

where *ŷ*_*i*_ is the predicted diffusion coefficient of the *i-*th origami, *y*_*i*_ is its experimental value, and *n* represents the number of considered DNA origamis. The lower the RMSE, the lower the differences between predicted and experimental values across samples (and hence, the better the model).

#### Hyperparameters optimization

Hyperparameters of each individual ML approach were optimized via grid search, as follows:

- Random Forest Regressor: “n_estimators” = [25, 50, 100, 200]; “max_depth” = [None, 10, 20, 30, 40]; “min_samples_split” = [2, 5, 10, 20]; “min_samples_leaf” = [1, 2, 4, 8].
- Support Vector Regressor: “C” = [0.1, 1.0, 10, 100]; “epsilon” = [0.01, 0.1, 0.5, 1.0]; “kernel” = [linear, poly, rbf, sigmoid]; “degree” = [2, 3, 4]; “gamma” = [scale, auto, 0.001, 0.01, 0.1, 1].
- Gaussian Process Regressor (k1): “kernel” = [RBF]; “length_scale” = [10, 5, 1.0, 0.1, 0.01, 0.001]; “noise_variance” = [1.0, 0.5, 0.25, 0.1]; “C” = [1] (constant value kernel magnitude): “alpha” = [1e-3, 1e-5, 1e-7, 1e-9].
- Gaussian Process Regressor (k2): “kernel” = [RBF]; “length_scale_1” = [10, 5, 1.0, 0.1]; “length_scale_2” = [0.01, 0.001, 0.0001]; “noise_variance” = [1.0, 0.5, 0.25, 0.1]; “C” = [1.0] (constant value kernel magnitude): “alpha” = [1e-3, 1e-5, 1e-7, 1e-9].
- XGBoost: “n_estimators” = [25, 50, 100, 200]; “max_depth” = [3, 6, 9, 12]; “learning_rate” = [0.4, 0.6, 0.8, 1.0]; “subsample” = [0.4, 0.6, 0.8, 1.0];“colsample_bytree” = [0.4, 0.6, 0.8, 1.0].

The hyperparameter combination yielding the lowest RMSE value (Eq. 4) in five-fold cross validation (on the training set) was chosen for follow-up validation.

#### Software and Code

All code was implemented in Python (v3.11.9). The machine learning models Random Forest Regressor, Support Vector Regressor, and Gaussian Process Regressor were implemented using the scikit-learn (v.1.5.0) ^55^. The XGBoost regressor model was implemented using the xgboost (v2.1.0) ^48^ package. Plots were made using combinations of the matplotlib (v3.8.4) ^56^ and seaborn (v0.13.2) ^57^ packages.

## Data availability

The raw data used for machine learning cycles, as well as the processed data used to create the manuscript’s figures, are available on GitHub at https://github.com/molML/DNA-origami-stability-prediction. The data at the time of publishing will be available at: XXX/zenodo.org/XXX.

## Code availability

The Python code to replicate and extend our model is openly accessible on GitHub at https://github.com/molML/DNA-origami-stability-prediction. The code at the time of publishing is available at: XXX/zenodo.XXX.

## CRediT author statement

JZA: Methodology (Experimental and Computational), Investigation, Software, Visualization, Validation, Formal analysis, Data Curation, Writing-Original Draft; AG: Methodology (Computational), Software, Validation, Formal analysis, Investigation, Writing-original Draft, Visualization. LP: Methodology, Investigation, Validation, Writing-Review & Editing. MTo: Methodology (Experimental), Investigation, Supervision. AA: Supervision, MTe: Supervision, FG: Conceptualization, Methodology, Resources, Writing-Original Draft, Supervision. TP: Conceptualization, Methodology, Resources, Writing-Original Draft, Supervision, Funding Acquisition.

## Acknowledgments

AG acknowledges support from the National Growth Fund Big Chemistry funded by the Dutch Ministry of Education, Culture and Science.

MTo thanks the Danish National Research Foundation (DNRF122) and Villum Fonden (Grant No. 9301) for intelligent drug delivery and sensing using microcontainers and nanomechanics (IDUN) and the Novo Nordisk Foundation (NNF17OC0026910).

JZA acknowledges the financial support provided by the Erasmus+ Traineeship Programme from the European Commission.

TP acknowledges funding from the Dutch Research Council (NWO) under the grant https://doi.org/10.61686/LVZRW92421.

We gratefully acknowledge the Institute for Complex Molecular Systems (ICMS) for providing laboratory facilities.

## Bibliography

1. Rothemund, P. W. K. Folding DNA to create nanoscale shapes and patterns. Nature 440, 297–302 (2006).

2. Douglas, S. M. et al. Self-assembly of DNA into nanoscale three-dimensional shapes. Nature 459, 414–418 (2009).

3. Dey, S. et al. DNA origami. Nat Rev Methods Primers 1, 13 (2021).

4. Jiang, Q., Shang, Y., Xie, Y. & Ding, B. DNA Origami: From Molecular Folding Art to Drug Delivery Technology. Adv Mater 36, 2301035 (2024).

5. Kretzmann, J. A. et al. Gene-encoding DNA origami for mammalian cell expression. Nat Commun 14, 1017 (2023).

6. Wu, X. et al. Genetically Encoded DNA Origami for Gene Therapy In Vivo. J Am Chem Soc 145, 9343–9353 (2023).

7. Li, J. et al. Tetrahedral DNA FrameworkLBased Spherical Nucleic Acids for Efficient siRNA Delivery. Angew Chem Int Ed 64, e202416988 (2025).

8. Comberlato, A., Koga, M. M., Nüssing, S., Parish, I. A. & Bastings, M. M. C. Spatially Controlled Activation of Toll-like Receptor 9 with DNA-Based Nanomaterials. Nano Lett 22, 2506–2513 (2022).

9. Knappe, G. A.Wamhoff, E.-C. & Bathe, M. Functionalizing DNA origami to investigate and interact with biological systems. Nat Rev Mater 8, 123–138 (2022).

10. Veneziano, R. et al. Role of nanoscale antigen organization on B-cell activation probed using DNA origami. Nat Nanotechnol 15, 716–723 (2020).

11. Berger, R. M. L. et al. Nanoscale FasL Organization on DNA Origami to Decipher Apoptosis Signal Activation in Cells. Small 17, 2101678 (2021).

12. Pfeiffer, M. et al. Single antibody detection in a DNA origami nanoantenna. iScience 24, 103072 (2021).

13. Wang, J. et al. Artificial molecular communication network based on DNA nanostructures recognition. Nat Commun 16, 244 (2025).

14. Ijäs, H. et al. DNA origami signal amplification in lateral flow immunoassays. Nat Commun 16, 3216 (2025).

15. Zhan, P. et al. Recent Advances in DNA Origami-Engineered Nanomaterials and Applications. Chem Rev 123, 3976–4050 (2023).

16. Linko, V. & Keller, A. Stability of DNA Origami Nanostructures in Physiological Media: The Role of Molecular Interactions. Small 19, 2301935 (2023).

17. Kim, H., Surwade, S. P., Powell, A., O’Donnell, C. & Liu, H. Stability of DNA origami nanostructure under diverse chemical environments. Chem Mater 26, 5265–5273 (2014).

18. Ramakrishnan, S., Ijäs, H., Linko, V. & Keller, A. Structural stability of DNA origami nanostructures under application-specific conditions. Comput Struct Biotechnol J 16, 342–349 (2018).

19. Hanke, M., Tomm, E., Grundmeier, G. & Keller, A. Effect of Ionic Strength on the Thermal Stability of DNA Origami Nanostructures. ChemBioChem 24, e202300338 (2023).

20. Chandrasekaran, A. R. Nuclease resistance of DNA nanostructures. Nat Rev Chem 5, 225–239 (2021).

21. Kemper, U., Weizenmann, N., Kielar, C., Erbe, A. & Seidel, R. Heavy Metal Stabilization of DNA Origami Nanostructures. Nano Lett 24, 2429–2436 (2024).

22. Rajendran, A., Krishnamurthy, K., Giridasappa, A., Nakata, E. & Morii, T. Stabilization and structural changes of 2D DNA origami by enzymatic ligation. Nucleic Acids Res 49, 7884– 7900 (2021).

23. Nguyen, M. K. et al. Ultrathin Silica Coating of DNA Origami Nanostructures. Chemistry of Materials 32, 6657–6665 (2020).

24. Gerling, T., Kube, M., Kick, B. & Dietz, H. Sequence-Programmable Covalent Bonding of Designed DNA Assemblies. Sci Adv 4, eaau1157 (2018).

25. Kiviaho, J. K. et al. Cationic polymers for DNA origami coating-examining their binding efficiency and tuning the enzymatic reaction rates. Nanoscale 8, 11674–11680 (2016).

26. Agarwal, N. P., Matthies, M., Gür, F. N., Osada, K. & Schmidt, T. L. Block Copolymer Micellization as a Protection Strategy for DNA Origami. Angew Chem Int Ed 129, 5552–5556 (2017).

27. Kopatz, I. et al. Packaging of DNA origami in viral capsids. Nanoscale 11, 10160– 10166 (2019).

28. Julin, S., Nonappa Shen, B., Linko, V. & Kostiainen, M. A. DNALOrigamiLTemplated Growth of Multilamellar Lipid Assemblies. Angew Chem Int Ed 60, 827–833 (2021).

29. Hannewald, N. et al. DNA Origami Meets Polymers: A Powerful Tool for the Design of Defined Nanostructures. Angew Chem Int Ed 60, 6218–6229 (2021).

30. Wassermann, L. M., Scheckenbach, M., Baptist, A. V., Glembockyte, V. & Heuer□ungemann, A. Full SiteLSpecific Addressability in DNA OrigamiLTemplated Silica Nanostructures. Adv Mater 35, 2212024 (2023).

31. Dou, B. et al. Machine Learning Methods for Small Data Challenges in Molecular Science. Chem Rev 123, 8736–8780 (2023).

32. Benjaminson, E., Taylor, R. E. & Travers, M. Predicting Nanorobot Shapes via Generative Models. arXiv:2101.12719 (2021).

33. Buterez, D. Scaling up DNA digital data storage by efficiently predicting DNA hybridisation using deep learning. Sci Rep 11, 20517 (2021).

34. Chiriboga, M. et al. Rapid DNA origami nanostructure detection and classification using the YOLOv5 deep convolutional neural network. Sci Rep 12, 3871 (2022).

35. Wang, Y., Jin, X. & Castro, C. Accelerating the characterization of dynamic DNA origami devices with deep neural networks. Sci Rep 13, 15196 (2023).

36. Cagirici, H. B., Budak, H. & Sen, T. Z. G4Boost: a machine learning-based tool for quadruplex identification and stability prediction. BMC Bioinformatics 23, 240 (2022).

37. Wang, D. et al. Functionalizing DNA nanostructures with natural cationic amino acids. Bioact Mater 6, 2946–2955 (2021).

38. Hanke, M. et al. Superstructure-dependent stability of DNA origami nanostructures in the presence of chaotropic denaturants. Nanoscale 15, 16590–16600 (2023).

39. Chen, C. et al. DNA origami frame filled with two types of single-stranded tiles. Nanoscale 14, 5340–5346 (2022).

40. Weck, J. M. & Heuer-Jungemann, A. Fully addressable, designer superstructures assembled from a single modular DNA origami. Nat Commun 16, 1556 (2025)

41. Bednarz, A., Sønderskov, S. M., Dong, M. & Birkedal, V. Ion-mediated control of structural integrity and reconfigurability of DNA nanostructures. Nanoscale 15, 1317–1326 (2022).

42. Benson, E. et al. DNA rendering of polyhedral meshes at the nanoscale. Nature 523, 441–444 (2015).

43. Rosier, B. J. H. M. et al. Proximity-induced caspase-9 activation on a DNA origami-based synthetic apoptosome. Nat Catal 3, 295–306 (2020).

44. Shaw, A. et al. Spatial control of membrane receptor function using ligand nanocalipers. Nat Methods 11, 841–846 (2014).

45. Zhang, B. et al. Investigation of nano-rods fabricated by the DNA origami method using static and dynamic light scattering. Mol Cryst and Liq Cryst 769, 1–9 (2025).

46. Ijäs, H., Hakaste, I., Shen, B., Kostiainen, M. A. & Linko, V. Reconfigurable DNA Origami Nanocapsule for pH-Controlled Encapsulation and Display of Cargo. ACS Nano 13, 5959–5967 (2019).

47. Breiman, L. Random Forests. Mach Learn 45, 5–32 (2001).

48. Chen, T. & Guestrin, C. XGBoost: a scalable tree boosting system. Proc. 22nd ACM SIGKDD Int Conf Knowl Discov Data Min 2016, 785–794 (2016).

49. Awad, M. & Khanna, R. Support vector regression. In Efficient Learning Machines: Theories, Concepts, and Applications for Engineers and System Designers (eds. Awad, M. & Khanna, R.) 67–80 (Apress, 2015).

50. Drucker, H., Burges, C. J. C., Kaufman, L., Smola, A. & Vapnik, V. Support vector regression machines. In Advances in Neural Information Processing Systems (eds. Mozer, M. C., Jordan, M. & Petsche, T.) 9 (MIT Press, 1996).

51. Rasmussen, C. E. & Williams, C. K. I. Gaussian Processes for Machine Learning (MIT Press, 2005).

52. Deringer, V. L. et al. Gaussian Process Regression for Materials and Molecules. Chem Rev 121, 10073–10141 (2021).

53. Ament, S. et al. Autonomous materials synthesis via hierarchical active learning of nonequilibrium phase diagrams. Sci Adv 7, eabg4930 (2025).

54. Stahl, E., Martin, T. G., Praetorius, F. & Dietz, H. Facile and scalable preparation of pure and dense DNA origami solutions. Angew Chem Int Ed 53, 12735–12740 (2014).

55. Pedregosa, F. et al. Scikit-learn: Machine Learning in Python. J Mach Learn Res 12, 2825–2830 (2011).

56. Hunter, J. D. Matplotlib: A 2D Graphics Environment. Comput Sci Eng 9, 90–95 (2007).

57. Waskom, M. Seaborn: statistical data visualization. J Open Source Softw 6, 3021 (2021).

