## Supporting Information for "Predicting DNA origami stability in physiological media by machine learning"

**SUPPLEMENTARY FIGURES**

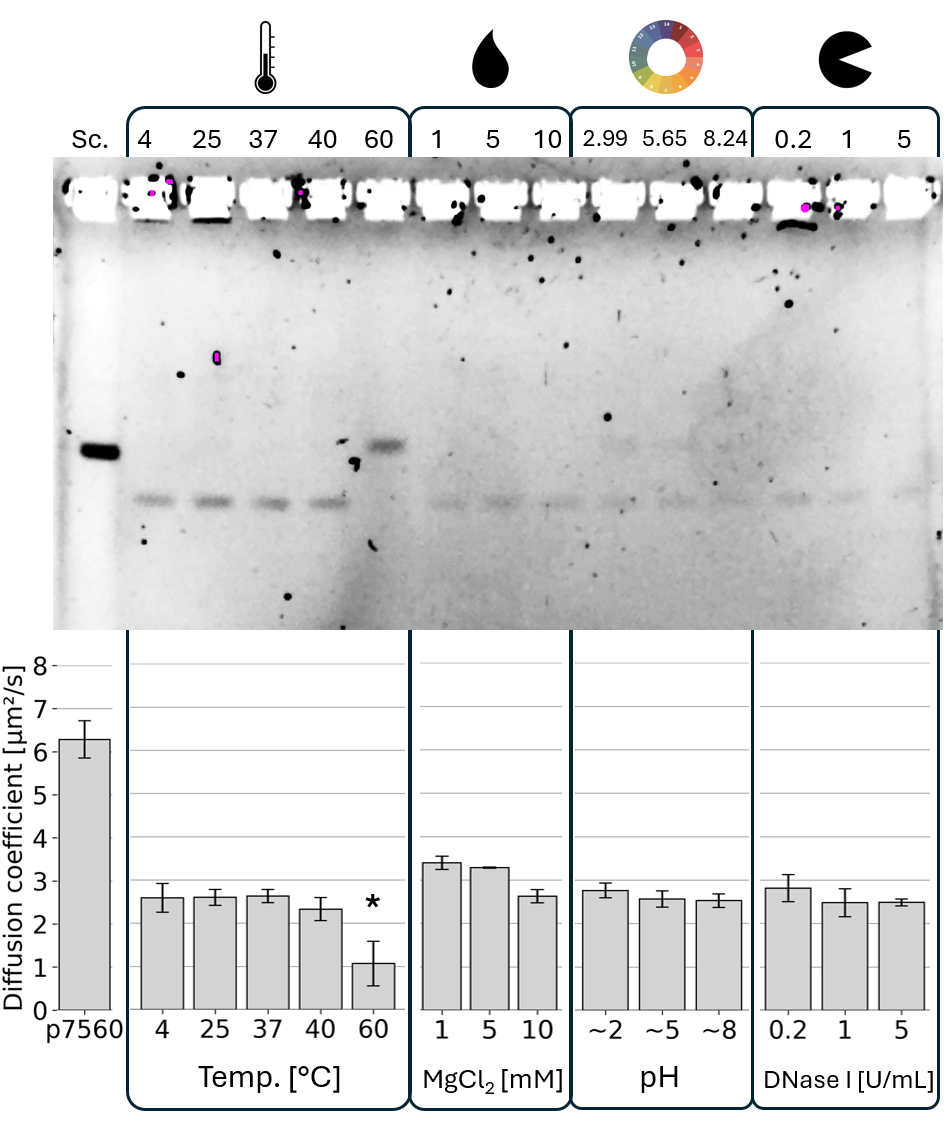

Figure S1. GE study of DNA origami rods after the exposure to different conditions for 60 min. Pink areas in the gel are artifacts introduced by the image acquisition device.

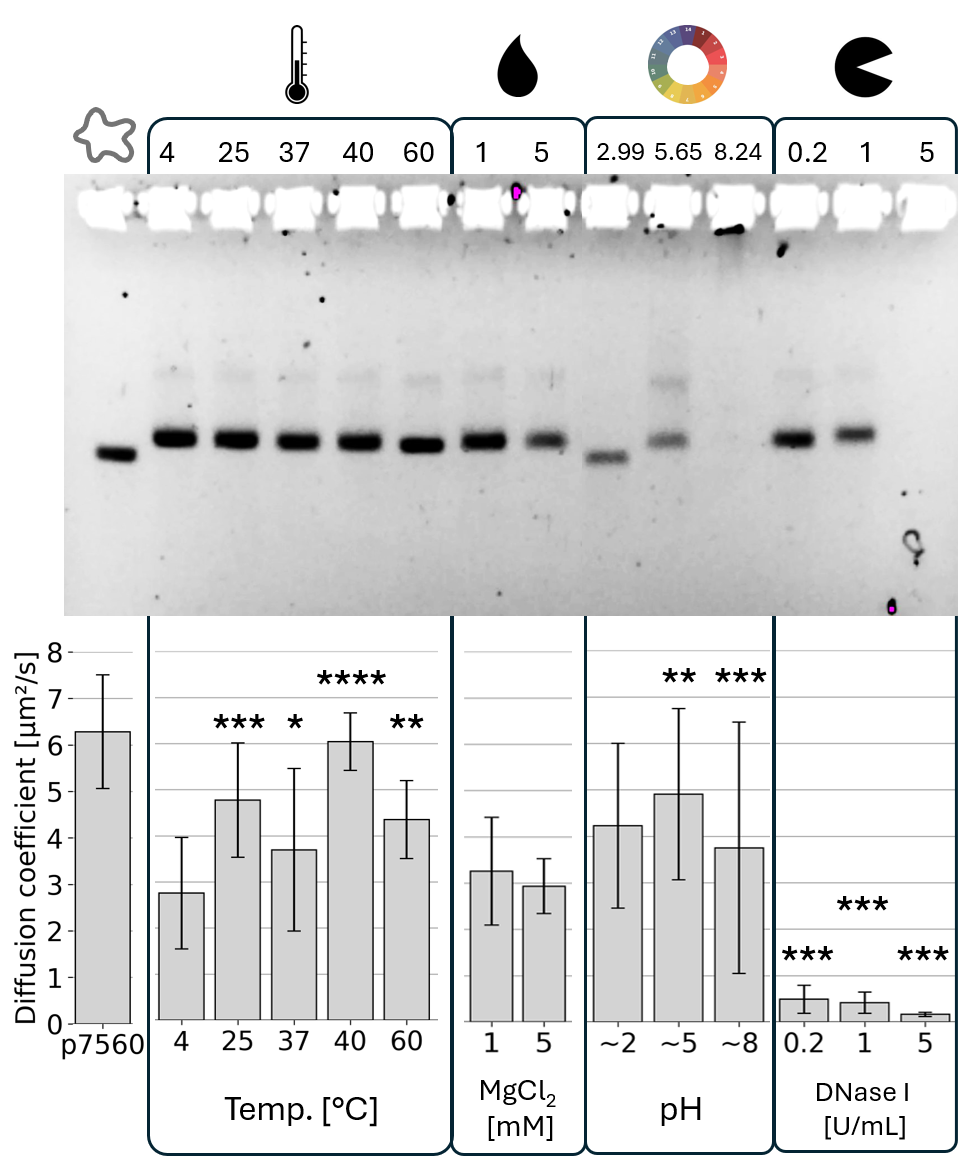

Figure S2. GE study of DNA origami icosahedrons after the exposure to different conditions for 60 min. Pink areas in the gel are artifacts introduced by the image acquisition device.

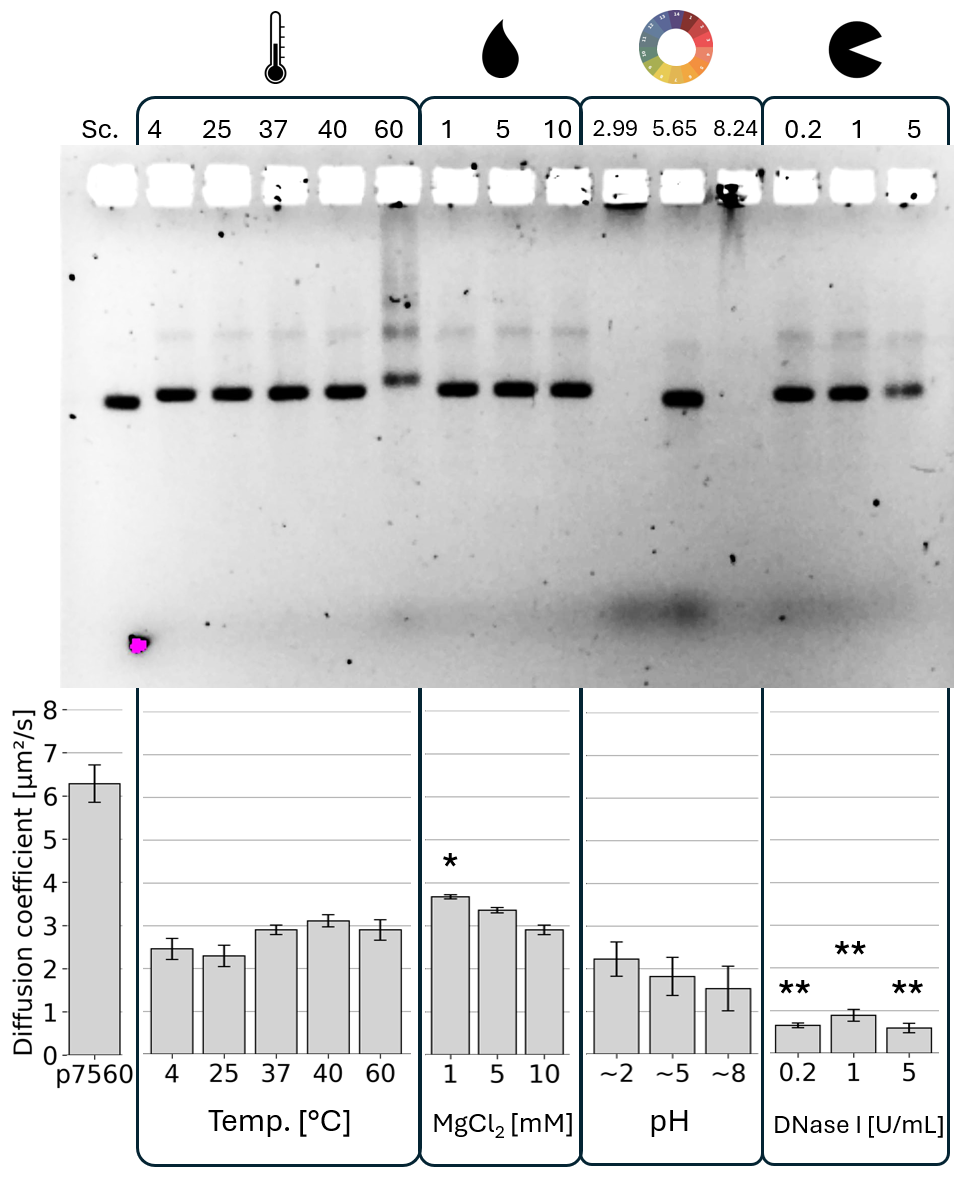

Figure S3. GE study of DNA origami rectangles after the exposure to different conditions for 60 min. Pink areas in the gel are artifacts introduced by the image acquisition device.

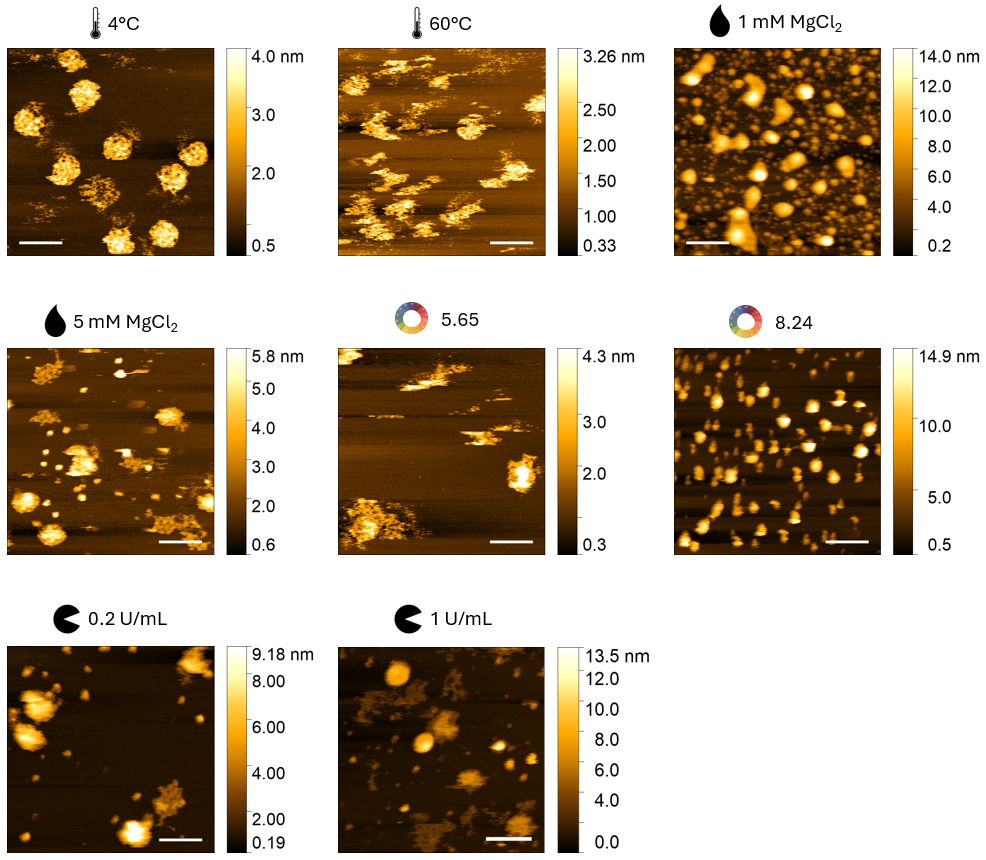

Figure S4. AFM height images of DNA origami icosahedrons corresponding to different unstable conditions acquired via tapping mode in liquid (scale bar = 100 nm).

**SUPPLEMENTARY TABLES**

Table S1. Staple strand sequences for the DNA origami rods. Location 5’ and Location 3’ refer to their corresponding positions in the caDNAno file.

| **Staple ID** | **Location 5'** | **Location 3'** | **Sequence 5' to 3'** |
| --- | --- | --- | --- |
| 1 | 0[51] | 14[42] | AAGACACCGCCTAACTGGCGCGGTAAGCCAACAGAGAT |
| 2 | 0[93] | 14[94] | GGAGAAAAATAACAGTACTTGAAACAAG |
| 3 | 0[114] | 17[104] | GGTTGCTGAATGAATTACCTTTTTTAATGGACTAAAGC |
| 4 | 0[135] | 14[136] | CAACCCTCAAATTACATGTCAATAAGAA |
| 5 | 0[156] | 17[146] | TCGGTTGGCAAACATCAAGAAAACAAAATTATCAATAT |
| 6 | 0[177] | 14[178] | TGTAGGAATTGCAAAAGCTTTTTATAGA |
| 7 | 0[198] | 17[188] | TGCAAAATATTTATTCATTTCAATTACCTGAGAGGAAG |
| 8 | 0[219] | 14[220] | AGCACAACTAACAAAATAAATGCTCTGA |
| 9 | 0[240] | 17[230] | GCCTCAATAGGATTGCTTTGAATACCAAGTTATAGATT |
| 10 | 0[261] | 14[262] | AACGATTTAGGAAACAAAAACTTTAGCT |
| 11 | 0[282] | 17[272] | CTTACAAACAACAGTACCTTTTACATCGGGAAAGTATT |
| 12 | 0[303] | 14[304] | TTGATTAAATACGTCAGTAAATTTTAAA |
| 13 | 0[324] | 17[314] | CGGTTATTAATTGCGTAGATTTTCAGGTTTACCTTTGC |
| 14 | 0[345] | 14[346] | ATTGTAACATCGTAAAACGTTAAATTTC |
| 15 | 0[366] | 17[356] | GAAACAAAGAACCATATCAAAATTATTTGCATATCATT |
| 16 | 0[387] | 14[388] | CACGCGGAATATAATGGAGCCTGTGCCA |
| 17 | 0[408] | 17[398] | TCATGATTATTTGTTTGGATTATACTTCTGATATCATC |
| 18 | 0[419] | 3[412] | AGTGAGCCATACGAAACCGTGCATCTGCAATGGGA |
| 19 | 1[73] | 17[58] | TCAATTAAAGACGCTGAGTGTGAGTGAATAACCTTGCTTCTG |
| 20 | 2[34] | 4[35] | CCGATTTGAGAAAGGAAGGGAGCGCGTA |
| 21 | 2[55] | 2[56] | TCGGAACGAAAGGAGCGGGCGCTAGGAATGTAAAGCACTAAA |
| 22 | 2[76] | 4[77] | TTAGTGCAGGCTATCAGGTCATTTTTGA |
| 23 | 2[118] | 4[119] | CTTCTAAAATCGATGAACGGTCAACCGT |
| 24 | 2[160] | 4[161] | GGCCAGTGTACCCCGGTTGATGAGAAAG |
| 25 | 2[202] | 4[203] | GGTAACGTTGTATAAGCAAATATGCAAT |
| 26 | 2[244] | 4[245] | CCAGCTGTAAAATTCGCATTACAACGCA |
| 27 | 2[286] | 4[287] | CTGTTGGACCAATAGGAACGCCATTATG |
| 28 | 2[328] | 4[329] | CGGAAACCTGTAGCCAGCTTTATAAAGC |
| 29 | 2[370] | 4[371] | AAGATCGACCCGTCGGATTCTATAAATC |
| 30 | 3[413] | 6[413] | TAGGTCAAGGTGGCATCAATTCTGTTTAGCTATATACGAACT |
| 31 | 4[34] | 7[34] | ACCACCAGCGTACTATGGTTGAAACAGGAGGCCGAGAATCCT |
| 32 | 4[55] | 4[56] | GTAGCGGGAGCACGTATAACGTGCTTTCACGCGCTGGCAAGT |
| 33 | 4[76] | 6[77] | GAGATCTAGAAGCAAAGCGGAACCCTGA |
| 34 | 4[107] | 0[115] | AATACAAGAGGTGGTTGCCCGCTTCTAATCTATAG |
| 35 | 4[118] | 6[119] | TCTAGCTGAAAGACTTCAAATTTCAGAA |
| 36 | 4[149] | 0[157] | TCAATCATATGCCAAGCGATACCGACAGTGCGAAA |
| 37 | 4[160] | 6[161] | GCCGGAGAACCAGACCGGAAGGTCATAA |
| 38 | 4[191] | 0[199] | GTGAGGAAGACCAGGGTGTGGGCACGAATATGGTT |
| 39 | 4[202] | 6[203] | GCCTGAGTACCTTTAATTGCTAGTAAAA |
| 40 | 4[233] | 0[241] | TTTATTTTGTGCGAAAGACATAAATCATTTCCACC |
| 41 | 4[244] | 6[245] | AGGATAATGGCTTAGAGCTTAGCGAGAG |
| 42 | 4[275] | 0[283] | CTTTTTTTTAGAAGGGCATGAGTAAACAGGGTTTT |
| 43 | 4[286] | 6[287] | ACCCTGTATGTTTTAAATATGATCATAA |
| 44 | 4[317] | 0[325] | AAGGGCCTTCCAGGCAAATAAAGACGGAGGACGCG |
| 45 | 4[328] | 6[329] | CTCAGAGATTCCATATAACAGATAACGC |
| 46 | 4[359] | 0[367] | GCAAGTAACACACTCCATCATGGTCATAGCTTCGG |
| 47 | 4[370] | 6[371] | ATACAGGTTTAGTTTGACCATCAGTTGA |
| 48 | 4[401] | 0[409] | TAATGACCGTCAGTTTGACAATTCCACACAATAAC |
| 49 | 5[46] | 0[52] | GACTCACGCTAGAAAGCCCTAAAGTCAAGTTTTTTGGGGTCA |
| 50 | 5[88] | 0[94] | ATCGTAGCTATTGCCTGCGACTTAACAATGTCCCGCCAGTTT |
| 51 | 5[130] | 0[136] | CGTTGATATTAATCGTACCAGGGTCTCGCCCTGGAGTGAAAT |
| 52 | 5[172] | 0[178] | CTCAAAGGGTAATCAGACGTTGTACCATCTGTAAGCAAATCC |
| 53 | 5[214] | 0[220] | TTGATTTTAAATTTAAAAGGCGATTTGAATCGGCTGACTTGC |
| 54 | 5[256] | 0[262] | CTGTTTATTTAATTTTTTCTTCGCTCTGACCTCCTGGTGGGC |
| 55 | 5[298] | 0[304] | TAACCAAAAACATCAAATTCAGGCTACGTGGTGCTTGTCGTA |
| 56 | 5[340] | 0[346] | TTCTTAAGCACATCAACACCGCTTGTACCGAGCTCGAACTGC |
| 57 | 5[382] | 0[388] | TACACATCCACCGTGGGGTATCGGGTGTGAAATTGTTACGCT |
| 58 | 6[55] | 6[56] | TCAGAGCTAATCAGTGAGGGCCTGATAAAGTCCTCGTTAGAA |
| 59 | 6[76] | 9[69] | CTATTATATTGTGTGGAGATTTGTATCATAGTACC |
| 60 | 6[118] | 9[111] | AACGAGAAACGAGGTTGACCCCCAGCGACCTCAGA |
| 61 | 6[160] | 9[153] | ATATTCAAACTTTGCTAAAACGAAAGAGTAGGAAC |
| 62 | 6[202] | 9[195] | TGTTTAGATAGGCTTTAAACGGGTAAAAAAACTAC |
| 63 | 6[244] | 9[237] | GCTTTTGCGGATATGAGGCTTTGAGGACTAGCGTA |
| 64 | 6[286] | 9[279] | CCCTCGTCAGTGAAGCAGCGAAAGACAGTAAATGA |
| 65 | 6[328] | 9[321] | CAAAAGGAGTAAATTGCAGGGAGTTAAACAGTTTC |
| 66 | 6[370] | 9[363] | GATTTAGTGTGAATACCATCGCCCACGCTTGCGAA |
| 67 | 6[412] | 9[405] | AACGGAACAGTCAGTTAAACAGCTTGATAGGCTCC |
| 68 | 9[49] | 9[48] | GCCTGAGTAGAATTTCCCACCGAGTAAAAGAGTCTATCACTT |
| 69 | 9[70] | 12[70] | GCCACCCCACCGTACTCAGGATTAAAGGTGAATTAATTGACG |
| 70 | 9[91] | 11[90] | CTCAGAATATAGCCCGGAATACACCGAC |
| 71 | 9[102] | 4[108] | CACTTATACCTGTTACTTAGCCGGATGACCAGAAGCCCGATA |
| 72 | 9[112] | 12[112] | GCCACCACGTCGAGAGGGTTGAATTAGAGCCAGCACAAAAGG |
| 73 | 9[133] | 11[132] | GATAGCACAGTACCAGGCGGACAGTAGC |
| 74 | 9[144] | 4[150] | CAAGCAAAAGGAACCGAACTGACCTTGAATCCAAAGCGACAG |
| 75 | 9[154] | 12[154] | CCATGTATAGGATTAGCGGGGCAAGGCCGGAAACGCACAATC |
| 76 | 9[175] | 11[174] | TTCGTCAGAGACTCCTCAAGATGAAACC |
| 77 | 9[186] | 4[192] | TACTACGTAATACAGACCAGGCGCACTGGATTAGAGAGTAAT |
| 78 | 9[196] | 12[196] | AACGCCTACATGAAAGTATTACGTAATCAGTAGCGATAAAAG |
| 79 | 9[217] | 11[216] | ACAGCCCTTCGGAACCTATTACAAGTTT |
| 80 | 9[228] | 4[234] | AGTTAAAGACATCTTGACAAGAACCAAAAGATTGCGGAAAAT |
| 81 | 9[238] | 12[238] | ACGATCTCAGTTAATGCCCCCACTGTAGCGCGTTTTGTTAGC |
| 82 | 9[259] | 11[258] | TTTCCAGGAGTAACAGTGCCCCATTTTC |
| 83 | 9[270] | 4[276] | TAGCATCGGACAAAGCTGCTCATTTTACCAGGCTCAACAATA |
| 84 | 9[280] | 12[280] | ATTTTCTTTTTAACGGGGTCATATTAGCGTTTGCCCCCAAAA |
| 85 | 9[301] | 11[300] | TAAACAACAGGAGTGTACTGGCATAATC |
| 86 | 9[312] | 4[318] | CAAGGCCGCTACACCAGAACGAGTAATTACGAAGTTTCCATA |
| 87 | 9[322] | 12[322] | AGCGGAGCATACATGGCTTTTCAGAGCCACCACCGGTTACCA |
| 88 | 9[343] | 11[342] | AACAACTATTTACCGTTCCAGCTCCCTC |
| 89 | 9[354] | 4[360] | GAAATAACCGTCAACTTTAATCATGAATACCGAGTAGACAAG |
| 90 | 9[364] | 12[364] | TAATAATAATGGAAAGCGCAGCAGAACCGCCACCCCCGAAGC |
| 91 | 9[385] | 11[384] | AATCTCCATAAATCCTCATTACACCACC |
| 92 | 9[396] | 4[402] | AAAACCGATACTGGCTCATTATACCAACATTCAATAACCTAC |
| 93 | 9[406] | 12[406] | AAAAGGAGGCCTTGATATTCACAGAACCACCACCAATAATAA |
| 94 | 10[37] | 5[45] | GGTTAATAACGTCCATCGAGAAGTGTTTTTAGGGAGCTCTTT |
| 95 | 10[79] | 5[87] | TATTCAGAACGTACAACCGAAATCCGCGACCGGTCTTTTTGC |
| 96 | 10[121] | 5[129] | TGCCCCTCATCTCATCTCGCAGACGGTCAATTAAACAGATCG |
| 97 | 10[163] | 5[171] | GATCCGTAACCACCAACAAAGAGGACAGATGCGGAATCCAAA |
| 98 | 10[205] | 5[213] | GAAGTAGCATGTTTCCAGGCTGACCTTCATCGGGTAATCCTT |
| 99 | 10[247] | 5[255] | AAAAAAGTTTGGCTACATCATTACCCAAATCCAAAATAATTG |
| 100 | 10[289] | 5[297] | AAGGTATGGGACCCTCATAAGGCTTGCCCTGCAACACTCAAC |
| 101 | 10[331] | 5[339] | CGTTGAGAATTGAGGCTTGGGCTTGAGATGGCAGATACTTGA |
| 102 | 10[373] | 5[381] | CAGTTTTTCAGACAACATACCTTATGCGATTGATTCATTAGA |
| 103 | 11[49] | 11[48] | GAAATACCTACACAGGGAACTCAAACTATCGGCCTGCTCATG |
| 104 | 11[91] | 14[84] | TTGAGCCATTGAGGGAGGGAACCGGTATTTTTTAT |
| 105 | 11[133] | 14[126] | ACCATTAATGGTTTACCAGCGCTTGCGGGTATTAA |
| 106 | 11[175] | 14[168] | ATCGATACACGGAATAAGTTTTTTTGCACAATCAA |
| 107 | 11[217] | 14[210] | GCCTTTACATACATAAAGGTGTAACGAGAGAAAAA |
| 108 | 11[259] | 14[252] | GGTCATAAAGACTCCTTATTAAAACAGCGCAGAAC |
| 109 | 11[301] | 14[294] | AAAATCAAAACGCAATAATAATTTTGTTATTCTGT |
| 110 | 11[343] | 14[336] | AGAGCCGGTAAGCAGATAGCCAGAATAACCAGTAA |
| 111 | 11[385] | 14[378] | CTCAGAGGAAATAGCAATAGCAACTGAATTAACAA |
| 112 | 12[69] | 15[73] | GAAATTAAATCAGATCATTACCGCGCCCAAATCGTCGCTATTAATT |
| 113 | 12[111] | 15[115] | GCGACATGCGTTTTTCATCGAGAACAAGACATAGCGATAGCTTAGA |
| 114 | 12[153] | 15[157] | AATAGAATTAAATCTCCTTATCATTCCAGTGAATTTATCAAAATCA |
| 115 | 12[195] | 15[199] | AAACGCATATCCTGAATTTACGAGCATGACCTCCGGCTTAGGTTGG |
| 116 | 12[237] | 15[241] | AAACGTACTAATTTACAATAGATAAGTCGATGCAAATCCAATCGCA |
| 117 | 12[279] | 15[283] | GAACTGGCAATCCATAAACAACATGTTCTTCAAATATATTTTAGTT |
| 118 | 12[321] | 15[325] | GAAGGAAAAAATAGGTACCGACAAAAGGAATGGTTTGAAATACCGA |
| 119 | 12[363] | 15[367] | CCTTTTTAGCGCATTTAGGCAGAGGCATTAAGAATAAACACCGGAA |
| 120 | 12[405] | 15[409] | GAGCAAGTCAGAGGCTTAATTGAGAATCTTAGTATCATATGCGTTA |
| 121 | 13[49] | 13[48] | TTCTGTAGCAAGCATTTTTTGACGCTCAATCGTCTAGGGACA |
| 122 | 14[41] | 17[41] | AGAACCCTATTAGTCTTTAATATAGCCCTAAAACAGGCGGTC |
| 123 | 14[83] | 17[83] | TTTCATCTTTCCCTTAGAATCCATAAATCAATATAAGCCAGC |
| 124 | 14[93] | 9[101] | CCGTCTAAGAACGCGAGTCAACCGATTTGGGATATAAGCCGC |
| 125 | 14[125] | 17[125] | ACCAAGTAGACGCTGAGAAGATTAACAATTTCATTCCTCAAA |
| 126 | 14[135] | 9[143] | CGGGAGGTTTTGAAGCCAATTCATCCATTAGTTTTGCTAGCC |
| 127 | 14[167] | 17[167] | TAATCGGGTCTGAGAGACTACAAGATGATGAAACAAATCAAC |
| 128 | 14[177] | 9[185] | AACCCCAGCTACAATTTAAGACACGCAGCACAGAGGCTCCAG |
| 129 | 14[209] | 17[209] | TAATATCATATAACTATATGTCGCGCAGAGGCGAACTTTAGG |
| 130 | 14[219] | 9[227] | ACACGTCTTTCCAGAGCGAAAATAGCGTCAGTGCCTATTCAT |
| 131 | 14[251] | 17[251] | GCGCCTGCAAAGAACGCGAGATAACGGATTCGCCTATAATAC |
| 132 | 14[261] | 9[269] | AATCATATTATTTATCCCATGATTGCCCCCTGTGCCTTACGT |
| 133 | 14[293] | 17[293] | CCAGACGTTCATCTTCTGACCATGAATATACAGTAATTCGAC |
| 134 | 14[303] | 9[311] | GTATAACGTCAAAAATGACCGAGGCCGGAACGATGATACTTT |
| 135 | 14[335] | 17[335] | TAAGAGATGTGATAAATAAGGCAGAAATAAAGAAATTTTAAA |
| 136 | 14[345] | 9[353] | GAGCATAAAAACAGGGAAAGAAAACCACCCTTCTCTGAAAAG |
| 137 | 14[377] | 17[377] | CGCCAACTAATTACTAGAAAAAAGGGTTAGAACCTAACCACC |
| 138 | 14[387] | 9[395] | TATCACCCTGAACAAAGAAACAATCCGCCACCAAACAAAAAA |
| 139 | 15[32] | 10[38] | GGCTTCTGACCGACCAGTAATAAAGAAATGGGAAAAACTGCT |
| 140 | 15[74] | 10[80] | AATGTAGGAATATAGAAGGCTTATGGTAATTCACCGTGGTG |
| 141 | 15[116] | 10[122] | TTAACCGCACAGCGAACCTCCCGACCAAAGAAAATCACTAAG |
| 142 | 15[158] | 10[164] | TAGCTGTCTTAAGATTAGTTGCTAATTTTGTTCACCAAGAAG |
| 143 | 15[200] | 10[206] | GTTCCATCCTAATCTTACCAACGCGCAACATACAGAATTTCT |
| 144 | 15[242] | 10[248] | AGATTTATCAGCCAGTTACAAAATCGCAGTATCATCGGGTAT |
| 145 | 15[284] | 10[290] | AATACGACAAAATAAGAAACGATTCGGAATAATCTTTTTAAT |
| 146 | 15[326] | 10[332] | CCGATATAAACAGCCTTTACAGAGGAACAAAGAACCGCTAAG |
| 147 | 15[368] | 10[374] | TCAATGTAATTAGACGGGAGAATTTATCTTATCAGAGCAAGC |
| 148 | 15[410] | 10[416] | TACAGTAGGGGTAATTGAGCGCTAAAGCCCAGAGCCGCGACG |
| 149 | 17[42] | 2[35] | AGTATTAGGCGAAAAACCGTCACCCAAAGGAGCCC |
| 150 | 17[59] | 1[72] | CAACAGTGCCAACGTGGACTCCAACGTCGAGGTGCCGAATTG |
| 151 | 17[84] | 2[77] | AGCAAATACAAGAGTCCACTACCTTATGAGTGTCC |
| 152 | 17[126] | 2[119] | TATCAAAAAGAATAGCCCGAGATTTACGGGATGTT |
| 153 | 17[168] | 2[161] | AGTTGAATTGATGGTGGTTCCGGCCCTGAAACGAC |
| 154 | 17[189] | 2[182] | GTTATCTCCCAGCAGGCGAAACTCGTCGTTTCCCA |
| 155 | 17[210] | 2[203] | AGCACTAAAGCGGTCCACGCTAGGGGCCTAAGTTG |
| 156 | 17[252] | 2[245] | ATTTGAGAGCTGATTGCCCTTTCCGAACTATTACG |
| 157 | 17[273] | 2[266] | AGACTTTTTCACCAGTGAGACTGGTGTAGATCGGT |
| 158 | 17[294] | 2[287] | AACTCGTGGCGCCAGGGTGGTCTTAAGCTGCGCAA |
| 159 | 17[315] | 2[308] | CCGAACGGGAGAGGCGGTTTGTACCTCGAGCGCCA |
| 160 | 17[336] | 2[329] | AGTTTGAAATGAATCGGCCAATCCCCGGCTGGTGC |
| 161 | 17[357] | 2[350] | TTGCGGAACCTGTCGTGCCAGTTCGTAAGCCAGCT |
| 162 | 17[378] | 2[371] | AGAAGGATGCCCGCTTTCCAGGTTTCCTCCTCAGG |
| 163 | 1[15] | 0[15] | AAAACACTACGTGAACCATCTATCAGGGCGATGGCCAAAA |
| 164 | 3[8] | 2[8] | AAAAAGCCGGCGAACGTGGCAGAGCTTGACGGGGAAAAAA |
| 165 | 5[8] | 4[8] | AAAATGCGCCGCTACAGGGCCACCCGCCGCGCTTAAAAAA |
| 166 | 7[8] | 6[8] | AAAACAGGAACGGTACGCCATTAAAGGGATTTTAGAAAAA |
| 167 | 8[34] | 8[12] | ACGCAAATTAACCGTTGTAAAAA |
| 168 | 9[12] | 9[34] | AAAAGCAATACTTCTTTGATTAG |
| 169 | 10[34] | 10[12] | AATATCCAGAACAATATTAAAAA |
| 170 | 11[12] | 11[34] | AAAACCGCCAGCCATTGCAACAG |
| 171 | 12[34] | 12[12] | ATTATTTACATTGGCAGATAAAA |
| 172 | 13[12] | 14[5] | AAAATCACCAGTCACACTGAAAGCGTAAGAATACGAAAA |
| 173 | 15[5] | 15[31] | AAAATGGCACAGACAATATTTTTGAAT |
| 174 | 17[5] | 17[27] | AAAAACCAGCAGAAGATAAAACA |
| 175 | 17[28] | 16[5] | GAGGTGATCGCCATTAAAAATACCGAACGAACCAAAA |
| 176 | 0[442] | 17[432] | AAAAAAGCCTGGGGTGTCAAAA |
| 177 | 2[435] | 1[442] | AAAATGGGCGCATCGTGCCGGAAGCATAAAGTGTAAAAA |
| 178 | 4[435] | 3[435] | AAAACGCGAGCTGAAACGTTGGTGTAGAAAAA |
| 179 | 6[435] | 5[435] | AAAATACGTTAATAAATTTCATTTGGGGAAAA |
| 180 | 8[439] | 9[439] | AAAAAGCTTGCTTTCGAGGTATTGTATCGGTTTATCAAAA |
| 181 | 10[415] | 7[435] | ATTGCCTTTAGAATTTCGACGTTGGGAAGAAAAATCAAAA |
| 182 | 10[440] | 11[440] | AAAAAGGTTGAGGCAGGTCACGCCAGCATTGACAGGAAAA |
| 183 | 12[439] | 13[439] | AAAACCACAAGAATTGAGTTATATCAGAGAGATAACAAAA |
| 184 | 14[432] | 15[432] | AAAAAAAGCCAACGCTCAACAAATTCTTACCAGTATAAAA |
| 185 | 16[432] | 0[420] | AAAAATCAATATAATCCTGACAGATGATGGCAATCCTAATG |
| 186 | 2[97] | 4[98] | ATGCGCAAGAGTCTGGAGCAATAATGCC |
| 187 | 4[97] | 6[98] | GGAGAGGAAAAAGATTAAGAGTAAATCA |
| 188 | 6[97] | 9[90] | AAAATCATGCTCCAAAGCGCGAAACAAACGCCACC |
| 189 | 17[105] | 2[98] | ATCACCTTGAGTGTTGTTCCAAAATAACTGAATTC |
| 190 | 2[139] | 4[140] | GGAGAAGAAACTAGCATGTCAAATCACC |
| 191 | 4[139] | 6[140] | ATCAATATTTAATTCGAGCTTCCCCTCA |
| 192 | 6[139] | 9[132] | AATGCTTCATAAGGAATACACTAAAACATTTCAGG |
| 193 | 17[147] | 2[140] | CTGGTCAGCAAAATCCCTTATACTCTATTTTCTCA |
| 194 | 2[181] | 4[182] | GTCACGAAAAGCCCCAAAAACTAGGTAA |
| 195 | 4[181] | 6[182] | AGATTCACAACAGGTCAGGATAGCGTCC |
| 196 | 6[181] | 9[174] | AATACTGAACGGTGTGCCACTACGAAGGACTGAGT |
| 197 | 2[223] | 4[224] | TGCTGCATTGTAAACGTTAATTAGAACC |
| 198 | 4[223] | 6[224] | CTCATATATAAGAGGTCATTTAGTTTTG |
| 199 | 6[223] | 9[216] | CCAGAGGAAGAGTATTTTTCATGAGGAATCCACAG |
| 200 | 17[231] | 2[224] | AGAGCCGTGGCCCTGAGAGAGGCATTTCGGGGATG |
| 201 | 2[265] | 4[266] | GCGGGCCGTTAAATCAGCTCATTGCGGG |
| 202 | 4[265] | 6[266] | AGAAGCCAATATAATGCTGTAACGACGA |
| 203 | 6[265] | 9[258] | TAAAAACAACGTAAACGAGGGTAGCAACTGTCGTC |
| 204 | 2[307] | 4[308] | TTCGCCAAATAATTCGCGTCTCTAAATC |
| 205 | 4[307] | 6[308] | GGTTGTAAGTACGGTGTCTGGAGGCATA |
| 206 | 6[307] | 9[300] | GTAAGAGACGAGAATTTGCGGGATCGTCATTTTGC |
| 207 | 2[349] | 4[350] | TTCCGGCATTAAATGTGAGCGAAGAATT |
| 208 | 4[349] | 6[350] | AGCAAAACCAATTCTGCGAACACATTCA |
| 209 | 6[349] | 9[342] | ACTAATGTTTAATTATATATTCGGTCGCAGAAAGG |
| 210 | 2[391] | 4[392] | GACGACAAACAAACGGCGGATTAGTAGT |
| 211 | 4[391] | 6[392] | AGCATTAATTTCGCAAATGGTATTACAG |
| 212 | 6[391] | 9[384] | GTAGAAATTAAGAAGTTGCGCCGACAATCGTTGAA |
| 213 | 17[399] | 2[392] | ATATTCCCATTAATTGCGTTGTCCGCTCAGGGGAC |

Table S2. Staple strand sequences for the DNA origami icosahedrons.

| Staple ID | **Sequence 5' to 3'** |
| --- | --- |
| 1 | CCACCCTCAGAGCCACCACCCTCATAGCTATCTTACCGAAGCCCT |
| 2 | TTGCTTCTGTAAATCGTCGCTATTAATCAGAGCCGCCACCCTCAGAACCG |
| 3 | AGCCACCACCGGAACCGCCTCCCACTATATGTAAATGCTGATGCAAATCC |
| 4 | TTTTAAGAAAAGTAAGCAGATATAATCAAAATCACCGGAACCAG |
| 5 | TTATTAGCGTTTGCCATCTTTTCATGTTAGCAAACGTAGAAAAT |
| 6 | AATCGCAAGACAAAGAACGCGAGAAATCGGCATTTTCGGTCATAGCCCCC |
| 7 | CGTCAGACTGTAGCGCGTTTTCAATCATATGCGTTATACAAATTCTTACC |
| 8 | ACATACATAAAGGTGGCAACACGACAGAATCAAGTTTGCCTTTAG |
| 9 | GATAGCAGCACCGTAATCAGTAGAAAGGTGAATTATCACCGTCACCG |
| 10 | AGTATAAAGCCAACGCTCAACAGTAGGGAAACGTCACCAATGAAACCATC |
| 11 | ACCATTACCATTAGCAAGGCCATGTTCAGCTAATGCAGAACGCGC |
| 12 | ATCCTAATTTACGAGCATGTAGAGCCAGCAAAATCACCAGTAGC |
| 13 | ACTTGAGCCATTTGGGAATTAGATTTCATCGTAGGAATCATTAC |
| 14 | AATATTGACGGAAATTATTCATTTATAAAAGAAACGCAAAGACACCA |
| 15 | CGCGCCCAATAGCAAGCAAATCAACCGATTGAGGGAGGGAAGGTA |
| 16 | AGCGCCAAAGACAAAAGGGCGACATTCGAGCGTCTTTCCAGAGCCTAATT |
| 17 | AGAAACGATTTTTTGTTTAACGTATCAATAGAAAATTCATATGGTTTACC |
| 18 | CGGAATAAGTTTATTTTGTCACAGGAATACCCAAAAGAACTGGCATG |
| 19 | ATTAAGACTCCTTATTACGCAGTAGCCGAACAAAGTTACCAGAAGGA |
| 20 | AACCGAGGAAACGCAATAATAACAATTGAGTTAAGCCCAATAATAAG |
| 21 | AGCAAGAAACAATGAAATAGCAATACCGTTCCAGTAAGCGTCATACATGG |
| 22 | ATATCAGAGAGATAACCCACAAGCAAAAATGAAAATAGCAGCCTTTA |
| 23 | CTTTTGATGATACAGGAGTGTACTGGAAGTCAGAGGGTAATTGAGCGCTA |
| 24 | ATTAACTGAACACCCTGAACAGATTAGCGGGGTTTTGCTCAGTAC |
| 25 | ATAGGTGTATCACCGTACTCAGGGGAAGCGCATTAGACGGGAGA |
| 26 | CAGAGAGAATAACATAAAAACAGATATTATTTATCCCAATCCAAATA |
| 27 | TGCCAGTTACAAAATAAACAGCCAGGTTTAGTACCGCCACCCTC |
| 28 | ATCCTGAATCTTACCAACGCTAACAGATATAGAAGGCTTATCCGGTA |
| 29 | AGAACCGCCACCCTCAGAACCTATTTTGCACCCAGCTACAATTTT |
| 30 | TGAAGCCTTAAATCAAGATTAGTTGCAGTTTTGTCGTCTTTCCAGACGTT |
| 31 | TTTCAGCGGAGTGAGAATAGAAAGCGAACCTCCCGACTTGCGGGAGGTTT |
| 32 | TTCTAAGAACGCGAGGCGTTTTAATTAAACCAAGTACCGCACTCATC |
| 33 | GAGAACAAGCAAGCCGTTTTTATAACCAATCAATAATCGGCTGTCTT |
| 34 | TCCTTATCATTCCAAGAACGGGTATAGTTGCGCCGACAATGACAACAACC |
| 35 | ATCGCCCACGCATAACCGATATATTCCCTGAACAAGAAAAATAATATCCC |
| 36 | CTGTTTATCAACAATAGATAAGTCCATTAAACGGGTAAAATACGTAATGC |
| 37 | TCCAGACGACGACAATAAACAACGGCTTAATTGAGAATCGCCATATT |
| 38 | CACTACGAAGGCACCAACCTAAAACGCCGACAAAAGGTAAAGTAATTCTG |
| 39 | GTAATAAGAGAATATAAAGTAACCTGTCGTGCCAGCTGCATT |
| 40 | AATGAATCGGCCAACGCGCGGGGCAGAGGCATTTTCGAGCCA |
| 41 | TAACAACGCCAACATGTAATTTATAAGAATAAACACCGGAATCATAA |
| 42 | TTACTAGAAAAAGCCTGTTTAGTACTTTTTCAAATATATTTTAGTTA |
| 43 | TGTGATAAATAAGGCGTTAAAGGAGAGGCGGTTTGCGTATTGGGC |
| 44 | GCCCTTCACCGCCTGGCCCTGAGAATGGTTTGAAATACCGACCG |
| 45 | ATTTCATCTTCTGACCTAAATTTGTCTGAGAGACTACCTTTTTAACC |
| 46 | TCCGGCTTAGGTTGGGTTATATATTAATTTTCCCTTAGAATCCTTGA |
| 47 | GTGAATTTATCAAAATCATAGAGAGTTGCAGCAAGCGGTCCACGC |
| 48 | TCGGCAAAATCCCTTATAAATCAAGACGCTGAGAAGAGTCAATA |
| 49 | AAACATAGCGATAGCTTAGATTACATTTAACAATTTCATTTGAATTA |
| 50 | GAGCCGCCGCCAGCATTGACAGGTAAATCAATATATGTGAGTGAATAACC |
| 51 | CCTTTTTTAATGGAAACAGTACAAGCAAAAGAAGATGATGAAACAAA |
| 52 | CATCAAGAAAACAAAATTAATTAAAAGAATAGCCCGAGATAGGG |
| 53 | GAATTATTCATTTCAATTACCTGATTGCGTAGATTTTCAGGTTTAAC |
| 54 | TTGAGTGTTGTTCCAGTTTGGCAAGTTACAAAATCGCGCAGAGGC |
| 55 | CGGATTCGCCTGATTGCTTTGAATACGGGAGCCCCCGATTTAGAGCTTGA |
| 56 | AGGAGCGGGCGCTAGGGCGCTGGTACCTTTTACATCGGGAGAAACAATAA |
| 57 | GTCAGATGAATATACAGTAACAGTCCTGATTGTTTGGATTATACTTC |
| 58 | ACGTAAAACAGAAATAAAGAAAGGTTGAGGCAGGTCAGACGA |
| 59 | TTGGCCTTGATATTCACAAACTACCATATCAAAATTATTTGC |
| 60 | TGAATAATGGAAGGGTTAGAACCAACAGTTAATGCCCCCTGCCT |
| 61 | ATGATGGCAATTCATCAATATAACAAGTGTAGCGGTCACGCTGCGCGTAA |
| 62 | ATTTCGGAACCTATTATTCTGTATCATCATATTCCTGATTATCAG |
| 63 | GAAACCACCAGAAGGAGCGGAATGAGGATTTAGAAGTATTAGACTTT |
| 64 | CCACCACACCCGCCGCGCTTAATGCGAACATTATCATTTTGCGGAACAAA |
| 65 | ATTAATTTTAAAAGTTTGAGTTGATAGCCCTAAAACATCGCC |
| 66 | ATTAAAAATACCGAACGAACCTAAATCCTTTGCCCGAACGTT |
| 67 | ACAAACAATTCGACAACTCGTATTGGCAAATCAACAGTTGAAAGGAA |
| 68 | TAGAGCCGTCAATAGATAATACATTTAAACATGAAAGTATTAAGAGGCTG |
| 69 | CAGGCGGATAAGTGCCGTCGAGACTTTAGGAGCACTAACAACTAATAGAT |
| 70 | TTGAGGAAGGTTATCTAAAATATTAGGAACCCATGTACCGTAACACTGAG |
| 71 | CCCTCAATCAATATCTGGTCAGTACCAGCAGAAGATAAAACAGAGGT |
| 72 | TTTCGTCACCAGTACAAACTACAACGCCTTGCTGAACCTCAAATATCAAA |
| 73 | CAGCAAATGAAAAATCTAAAGCATCATGGCTCATTATACCAGTCAGGACG |
| 74 | TATTACAGGTAGAAAGATTCATCCTGCAACAGTGCCACGCTGAGAGCCAG |
| 75 | GAGGCGGTCAGTATTAACACCGCGGCACAGACAATATTTTTGAATGG |
| 76 | CTATTAGTCTTTAATGCGCGAACAAAGGGATTTTAGACAGGAACGGTACG |
| 77 | GACCTGAAAGCGTAAGAATACGTAGTTGAGATTTAGGAATACCA |
| 78 | CCAGAATCCTGAGAAGTGTTTTTATAGGCCAACAGAGATAGAACCCTTCT |
| 79 | ACCAGTAATAAAAGGGACATTCTACAATATTACCGCCAGCCATTGCA |
| 80 | CATTCAACTAATGCAGATACAATTGGCAGATTCACCAGTCACACG |
| 81 | CGTCTGAAATGGATTATTTACGCGTCCAATACTGCGGAATCG |
| 82 | TCATAAATATTCATTGAATCCACCTACATTTTGACGCTCAAT |
| 83 | ACAGGAAAAACGCTCATGGAAATTTGCGGATGGCTTAGAGCTTAATTGCT |
| 84 | CGGCCTTGCTGGTAATATCCAGAATCAGTGAGGCCACCGAGTAAAAG |
| 85 | GAATATAATGCTGTAGCTCAACATGTCCTGAGTAGAAGAACTCAAACTAT |
| 87 | GATTAGTAATAACATCACTTGGTTTTTTGGGGTCGAGGTGCCGTA |
| 88 | CGGGGAAAGCCGGCGAACGTGGCCCGTTGTAGCAATACTTCTTT |
| 89 | AGTCTGTCCATCACGCAAATTAATGCTTTCCTCGTTAGAATCAGAGC |
| 90 | GGGAGCTAAACAGGAGGCCGATTCCGCTACAGGGCGCGTACTATGGT |
| 91 | TGCTTTGACGAGCACGTATAACGGAGAAAGGAAGGGAAGAAAGCGAA |
| 92 | AAGCACTAAATCGGAACCCTAAAAACAAGAGTCCACTATTAAAGAAC |
| 93 | ACTACGTGAACCATCACCCAAATCAATTTAAATATGCAACTAAAGTACGG |
| 94 | ACGAGTAGATTTAGTTTGACCATAAAAACCGTCTATCAGGGCGATGGCCC |
| 95 | GTGGACTCCAACGTCAAAGGGCGTCCTGTTTGATGGTGGTTCCGAAA |
| 96 | TGGTTTGCCCCAGCAGGCGAAAATAGATACATTTCGCAAATGGTCAATAA |
| 97 | CCTGTTTAGCTATATTTTCATTTGGGCAGTGAGACGGGCAACAGCTGATT |
| 98 | GCCAGGGTGGTTTTTCTTTTCACACTCACATTAATTGCGTTGCGCTC |
| 99 | ACTGCCCGCTTTCCAGTCGGGAACTGATAAATTGTGTCGAAATCCGCGAC |
| 100 | TGGGGTGCCTAATGAGTGAGCTAGCGCGAGCTGAAAAGGTGGCATCA |
| 101 | CTGCTCCATGTTACTTAGCCGGAACGGCCGGAAGCATAAAGTGTAAAGCC |
| 102 | ACAATTCCACACAACATACGACAAAAAGATTAAGAGGAAGCCCGA |
| 103 | CCGGAAGCAAACTCCAACAGGTCGTGTGAAATTGTTATCCGCTC |
| 104 | ATTCTACTAATAGTAGTAGCATTAACAGTTGATTCCCAATTCTGCGA |
| 105 | TGTCTGGAAGTTTCATTCCATATAGGATTAGAGAGTACCTTTAATTG |
| 106 | CTCCTTTTGATAAGAGGTCATTTATTCGAGCTTCAAAGCGAACCAGA |
| 107 | AAGACTTCAAATATCGCGTTTTACCCTCAAATGCTTTAAACAGTTCA |
| 108 | ATAGTCAGAAGCAAAGCGGATTGCATAGGCGCAGACGGTCAATCATAAGG |
| 109 | ACCAGGCGCATAGGCTGGCTGACAAAAATCAGGTCTTTACCCTGACTATT |
| 110 | GAAAACGAGAATGACCATAAATCAAGAAGTTTTGCCAGAGGGGGTAA |
| 111 | TAGTAAAATGTTTAGACTGGATATAACGCCAAAAGGAATTACGAGGC |
| 112 | ACCAAAATAGCGAGAGGCTTTTGCAACTTCATCAAGAGTAATCTTGACAA |
| 113 | AGTGAATAAGGCTTGCCCTGACGACCCTCGTTTACCAGACGACGATAAAA |
| 114 | ATAGTAAGAGCAACACTATCATATAAAACGAACTAACGGAACAACAT |
| 115 | TTGGGAAGAAAAATCTACGTTAAAGAAACACCAGAACGAGTAGTAAA |
| 116 | ACCTTATGCGATTTTAAGAACCCTGTAGCATTCCACAGACAGCCC |
| 117 | AGTAAATGAATTTTCTGTATGGGCAACTTTAATCATTGTGAATT |
| 118 | TTGGGCTTGAGATGGTTTAATTTCTCCAAAAAAAAGGCTCCAAA |
| 119 | AGGAGCCTTTAATTGTATCGGTCAACGTAACAAAGCTGCTCATTC |
| 120 | GAACCGGATATTCATTACCCAAACACCCTCAGCAGCGAAAGACA |
| 121 | GCATCGGAACGAGGGTAGCAAAGAGGACAGATGAACGGTGTACAG |
| 122 | GAACCGAACTGACCAACTTTGAAATTATACCAAGCGCGAAACAAAGT |
| 123 | ACAACGGAGATTTGTATCATCGCAAAGAGGCAAAAGAATACACTAAA |
| 124 | ACACTCATCTTTGACCCCCAGCGCGGCTACAGAGGCTTTGAGGACTA |
| 125 | AAGACTTTTTCATGAGGAAGTTTGGTCGCTGAGGCTTGCAGGGAGTT |
| 126 | GAACCGGATATTCATTACCCAAACACCCTCAGCAGCGAAAGACA |
| 127 | AAAGGCCGCTTTTGCGGGATCGTTTTATCAGCTTGCTTTCGAGGTGA |
| 128 | ATTTCTTAAACAGCTTGATACCGGGAACAACTAAAGGAATTGCGAAT |
| 129 | AATAATTTTTTCACGTTGAAAATATTTTGCTAAACAACTTTCAACAG |
| 130 | TCATAGTTAGCGTAACGATCTAAGCCACCCTCAGAGCCACCACCCTC |
| 131 | ATTTTCAGGGATAGCAAGCCCAAGGGTTGATATAAGTATAGCCCGGA |
| 132 | AGACTCCTCAAGAGAAGGATTAGTAATAAGTTTTAACGGGGTCAGTG |
| 133 | CCTTGAGTAACAGTGCCCGTATAAAATAAATCCTCATTAAAGCCAGA |
| 134 | ATGGAAAGCGCAGTCTCTGAATTGAGCCGCCACCAGAACCACCACCA |

Table S3. Staple strand sequences for the DNA origami rectangles. Location 5’ and Location 3’ refer to their corresponding positions in the caDNAno file.

| **Staple ID** | **Location 5'** | **Location 3'** | **Sequence 5' to 3'** |
| --- | --- | --- | --- |
| 1 | 0[79] | 1[63] | ACTGAGTTTCGTCACCAGTACAAATCATAGTT |
| 2 | 0[111] | 1[95] | GCAAGCCCAATAGGAACCCATGTACGTCTTTC |
| 3 | 0[143] | 1[127] | CCTCAGAGCCACCACCCTCATTTTGTATGGGA |
| 4 | 0[175] | 0[144] | CCCTCAGAACCGCCACCCTCAGAACCGCCAC |
| 5 | 0[207] | 1[191] | TATCACCGTACTCAGGAGGTTTAGATTATTCT |
| 6 | 0[239] | 1[223] | GGGTTGATATAAGTATAGCCCGGATGAGACTC |
| 7 | 1[64] | 3[63] | AGCGTAACAAAAGGCTCCAAAAGGTTCGAGGT |
| 8 | 1[96] | 3[95] | CAGACGTTAATAATTTTTTCACGTCGATAGTT |
| 9 | 1[128] | 3[127] | TTTTGCTAAATAGAAAGGAACAACGCCCACGC |
| 10 | 1[160] | 2[144] | TGCCCCCTAACAGTGCCCGTATAATTTCAGC |
| 11 | 1[192] | 3[191] | GAAACATGTAATAAGTTTTAACGGAGGTTGAG |
| 12 | 1[224] | 3[223] | CTCAAGAGCATGGCTTTTGATGATTATTCACA |
| 13 | 2[79] | 0[80] | CTCCAAAAGATCTAAAGTTTTGTCCGTAAC |
| 14 | 2[111] | 0[112] | TTGCGAATAGTAAATGAATTTTCTCAGGGATA |
| 15 | 2[143] | 1[159] | GGAGTGAGAACAACTTTCAACAGACAGTTAA |
| 16 | 2[175] | 0[176] | CCTTGAGTGCCTATTTCGGAACCTTACCGCCA |
| 17 | 2[207] | 0[208] | TGTACTGGAAAGTATTAAGAGGCATAGGTG |
| 18 | 2[239] | 0[240] | GCGTCATAAAGGATTAGGATTAGCCGTCGAGA |
| 19 | 3[64] | 5[63] | GAATTTCTCAACGGCTACAGAGGCTTCCATTA |
| 20 | 3[96] | 5[95] | GCGCCGACGCAGCGAAAGACAGCACTACGAAG |
| 21 | 3[128] | 5[127] | ATAACCGATAAAGGCCGCTTTTGCAAAAGAAT |
| 22 | 3[160] | 4[144] | AGAGCCGCAGAGCCGCCACCAGAAAGGCTTG |
| 23 | 3[192] | 5[191] | GCAGGTCATCAGAACCGCCACCCTTTTGCCTT |
| 24 | 3[224] | 5[223] | AACAAATAACCGGAACCGCCTCCCTCATCGGC |
| 25 | 4[79] | 2[80] | GAGGGTAGTAAACAGCTTGATACTGAAAAT |
| 26 | 4[111] | 2[112] | CACCCTCAAATGACAACAACCATCTAAAGGAA |
| 27 | 4[143] | 3[159] | CAGGGAGTTATATTCGGTCGCTGCCACCACC |
| 28 | 4[175] | 2[176] | CCACCCTCCGCCAGCATTGACAGGGGTCAGTG |
| 29 | 4[207] | 2[208] | CGCCACCCGACGATTGGCCTTGAACAGGAG |
| 30 | 4[239] | 2[240] | GAGCCACCAATCCTCATTAAAGCCTCCAGTAA |
| 31 | 5[64] | 7[63] | AACGGGTACGACCTGCTCCATGTTATAAGGGA |
| 32 | 5[96] | 7[95] | GCACCAACATCATCGCCTGATAAAAGGACAGA |
| 33 | 5[128] | 7[127] | ACACTAAACAAGCGCGAAACAAAGTAGGCTGG |
| 34 | 5[160] | 6[144] | GTAATCAGAATGAAACCATCGATACCCCAGC |
| 35 | 5[192] | 7[191] | TAGCGTCACCATTACCATTAGCAAAGCGCCAA |
| 36 | 5[224] | 7[223] | ATTTTCGGGGAATTAGAGCCAGCAGATTGAGG |
| 37 | 6[79] | 4[80] | GAAATCCGAAATACGTAATGCCATCGGAAC |
| 38 | 6[111] | 4[112] | AGATTTGTCTAAAACGAAAGAGGCGGGATCGT |
| 39 | 6[143] | 5[159] | GATTATACACACTCATCTTTGACGCAGCACC |
| 40 | 6[175] | 4[176] | ACGTCACCTAGCGACAGAATCAAGCAGAGCCA |
| 41 | 6[207] | 4[208] | CAGTAGCAGACTGTAGCGCGTTTTCAGAGC |
| 42 | 6[239] | 4[240] | GCCATTTGTCATAGCCCCCTTATTCGGAACCA |
| 43 | 7[64] | 9[63] | ACCGAACTAAACACCAGAACGAGTCTTTAATC |
| 44 | 7[96] | 9[95] | TGAACGGTTCATTCAGTGAATAAGTTAAGAAC |
| 45 | 7[128] | 9[127] | CTGACCTTATTCATTACCCAAATCTTGGGAAG |
| 46 | 7[160] | 8[144] | AATAGAAAGAATAAGTTTATTTTGTGACAAG |
| 47 | 7[224] | 9[223] | GAGGGAAGAGCAAACGTAGAAAATAAGGAAAC |
| 48 | 8[79] | 6[80] | CTGACGAGGACCAACTTTGAAAGTTGTGTC |
| 49 | 8[111] | 6[112] | AAAGCTGCGTACAGACCAGGCGCATACAACGG |
| 50 | 8[143] | 7[159] | AACCGGATCATCAAGAGTAATCTTCACAATC |
| 51 | 8[175] | 6[176] | ACACCACGATTCATATGGTTTACCGGCCGGAA |
| 52 | 8[207] | 6[208] | TAAAGGTGGGGCGACATTCAACCAAATCAC |
| 53 | 8[239] | 6[240] | AGTATGTTGTAAATATTGACGGAACGACTTGA |
| 54 | 9[64] | 11[63] | ATTGTGAAACATAACGCCAAAAGGTAACCCTC |
| 55 | 9[96] | 11[95] | TGGCTCATTTTAGGAATACCACATCAAAATAG |
| 56 | 9[128] | 11[127] | AAAAATCTTTATTACAGGTAGAAATTGCCAGA |
| 57 | 9[192] | 11[191] | AGATAGCCAAGAATTGAGTTAAGCGTTTAACG |
| 58 | 9[224] | 11[223] | CGAGGAAAATTGAGCGCTAATATCTACAGAGA |
| 59 | 10[79] | 8[80] | ATGCAGATTTACCTTATGCGATTGCTTGCC |
| 60 | 10[111] | 8[112] | AGTTGAGATATACCAGTCAGGACGAACGTAAC |
| 61 | 10[143] | 9[159] | GAACAACAACGTTAATAAAACGAAATAGCTA |
| 62 | 10[175] | 8[176] | AAGAGCAAAAGCCCTTTTTAAGAAACGCAAAG |
| 63 | 10[207] | 8[208] | TAACCCACGAACAAAGTTACCAGACATACA |
| 64 | 10[239] | 8[240] | AGAGGGTACGCAATAATAACGGAATATTACGC |
| 65 | 11[64] | 13[63] | GTTTACCAACGAGAATGACCATAAAGTCAGAA |
| 66 | 11[96] | 13[95] | CGAGAGGCTCCCCCTCAAATGCTTATTAAGAG |
| 67 | 11[128] | 13[127] | GGGGGTAAATACTGCGGAATCGTCGCGTTTTA |
| 68 | 11[160] | 12[144] | AATCCAAAATAAACAGCCATATTAACTGGAT |
| 69 | 11[192] | 13[191] | TCAAAAATGTCTTTCCAGAGCCTAAGGCTTAT |
| 70 | 11[224] | 13[223] | GAATAACATTTATCCTGAATCTTAGTTTTAGC |
| 71 | 12[79] | 10[80] | TTCAGAAAGACGACGATAAAAACTCAACTA |
| 72 | 12[111] | 10[112] | TCATTGAATTTTGCAAAAGAAGTTGATTCATC |
| 73 | 12[175] | 10[176] | GTTACAAATAAGAAACGATTTTTTCCAATAAT |
| 74 | 12[207] | 10[208] | TAACGAGCGAAAATAGCAGCCTTAGAGAGA |
| 75 | 12[239] | 10[240] | GCTACAATTAAAAACAGGGAAGCGACAAAGTC |
| 76 | 13[64] | 15[63] | GCAAAGCGTGGCTTAGAGCTTAATTAAATATG |
| 77 | 13[96] | 15[95] | GAAGCCCGGCTCCTTTTGATAAGAGTTTCATT |
| 78 | 13[128] | 15[127] | ATTCGAGCCAACAGGTCAGGATTACTGCGAAC |
| 79 | 13[160] | 14[144] | AATAGCAAATCGTAGGAATCATTAACCGGAA |
| 80 | 13[192] | 15[191] | CCGGTATTTCATCGAGAACAAGCAACGCGCCT |
| 81 | 13[224] | 15[223] | GAACCTCCCCAAGAACGGGTATTATGAACAAG |
| 82 | 14[79] | 12[80] | TTTGCGGAGATTGCATCAAAAAGTAAACAG |
| 83 | 14[111] | 12[112] | CTTTAATTAAAGACTTCAAATATCATAAATAT |
| 84 | 14[143] | 13[159] | GCAAACTCTTCAAAGCGAACCAGCCGCGCCC |
| 85 | 14[175] | 12[176] | TTATTTTCGCAAATCAGATATAGAATTTGCCA |
| 86 | 14[207] | 12[208] | TACCGCACCTAAGAACGCGAGGCCCAACGC |
| 87 | 14[239] | 12[240] | TTATCATTCGACTTGCGGGAGGTTTGCACCCA |
| 88 | 15[64] | 17[63] | CAACTAAACTACTAATAGTAGTAGAAAGAATT |
| 89 | 15[96] | 17[95] | CCATATAATGGGGCGCGAGCTGAACAGAGCAT |
| 90 | 15[128] | 17[127] | GAGTAGATTGGTCAATAACCTGTTACATTATG |
| 91 | 15[160] | 16[144] | AACAACATAATTCTGTCCAGACGAGATACAT |
| 92 | 15[192] | 17[191] | GTTTATCAAGAGAATATAAAGTACAAAAGCCT |
| 93 | 15[224] | 17[223] | AAAAATAATTTAGGCAGAGGCATTAAATTCTT |
| 94 | 16[79] | 14[80] | CATCAATTGTACGGTGTCTGGAAGGTCATT |
| 95 | 16[111] | 14[112] | TTTTCATTCAGTTGATTCCCAATTGAGAGTAC |
| 96 | 16[143] | 15[159] | TTCGCAAATTAGTTTGACCATTACGACAATA |
| 97 | 16[175] | 14[176] | GGTAAAGTGTTCAGCTAATGCAGAAGCCGTTT |
| 98 | 16[207] | 14[208] | CAGTAATAACAATAGATAAGTCCAACCAAG |
| 99 | 16[239] | 14[240] | ACATGTAATATCCCATCCTAATTTGTCTTTCC |
| 100 | 17[64] | 19[63] | AGCAAAATGTAAAGATTCAAAAGGCAATATGA |
| 101 | 17[96] | 19[95] | AAAGCTAAATTTTAAATGCAATGCATTAATGC |
| 102 | 17[128] | 19[127] | ACCCTGTACGCAAGGATAAAAATTGATCTACA |
| 103 | 17[160] | 18[144] | AACACCGGTAAATAAGGCGTTAAAAAGCCTT |
| 104 | 17[192] | 19[191] | GTTTAGTAAAATTTAATGGTTTGAGGTCTGAG |
| 105 | 17[224] | 19[223] | ACCAGTATATATATTTTAGTTAATTTAGGTTG |
| 106 | 18[79] | 16[80] | ATGTGTAGTAAGCAATAAAGCCTAAGGTGG |
| 107 | 18[111] | 16[112] | CCTCATATATCGGTTGTACCAAAATAGCTATA |
| 108 | 18[143] | 17[159] | TATTTCAAATACTTTTGCGGGAGTAAGAATA |
| 109 | 18[175] | 16[176] | CCGTGTGAAATCATAATTACTAGACGACAAAA |
| 110 | 18[207] | 16[208] | TCTGACCTTCATATGCGTTATACTTCGAGC |
| 111 | 18[239] | 16[240] | TTTTTCAAAAAGCCAACGCTCAACCAACGCCA |
| 112 | 19[64] | 21[63] | TATTCAACTCAGAAAAGCCCCAAATGTAAACG |
| 113 | 19[96] | 21[95] | CGGAGAGGGCATGTCAATCATATGTTAAATTT |
| 114 | 19[128] | 21[127] | AAGGCTATAAGAGAATCGATGAACCCAATAGG |
| 115 | 19[160] | 20[144] | AATAGTGATAGATTAAGACGCTGAGAGTCTG |
| 116 | 19[224] | 21[223] | GGTTATATCTTCTGTAAATCGTCGTTACATTT |
| 117 | 20[79] | 18[80] | GTTGATAACGTTCTAGCTGATAACTGAGTA |
| 118 | 20[111] | 18[112] | TAAAACTAGTAGCTATTTTTGAGATTTAGAAC |
| 119 | 20[143] | 19[159] | GAGCAAACCAGGTCATTGCCTGAGAAGAGTC |
| 120 | 20[175] | 18[176] | CGATAGCTATTTATCAAAATCATAAATACCGA |
| 121 | 20[207] | 18[208] | TTAATTTTTTTTTAACCTCCGGCTTCATCT |
| 122 | 20[239] | 18[240] | TAACCTTGAACTATATGTAAATGCGAGAAAAC |
| 123 | 21[64] | 23[63] | TTAATATTACGGCGGATTGACCGTCATCGTAA |
| 124 | 21[96] | 23[95] | TTGTTAAATAACAACCCGTCGGATGACGACGA |
| 125 | 21[128] | 23[127] | AACGCCATGCCAGCTTTCATCAACACTCCAGC |
| 126 | 21[160] | 22[144] | TCAATTACTCGCGCAGAGGCGAATTCTGGCC |
| 127 | 21[192] | 23[191] | AAACATCACGCCTGATTGCTTTGAATAATGGA |
| 128 | 21[224] | 23[223] | AACAATTTCCTTTTACATCGGGAGAATTATTT |
| 129 | 22[79] | 20[80] | GGGAACAATTGTTAAAATTCGCATACCCCG |
| 130 | 22[111] | 20[112] | TGAGCGAGTCAGCTCATTTTTTAAGGTAATCG |
| 131 | 22[143] | 21[159] | TTCCTGTACAAAAATAATTCGCGTATTCATT |
| 132 | 22[175] | 20[176] | TTACAAAACTGAGCAAAAGAAGATAAACATAG |
| 133 | 22[207] | 20[208] | AACGGATTAGAAAACAAAATTAACTATTAA |
| 134 | 22[239] | 20[240] | TAACAGTACATTTGAATTACCTTTTGAGTGAA |
| 135 | 23[64] | 25[63] | CCGTGCATGCTATTACGCCAGCTGTTGGGTAA |
| 136 | 23[96] | 25[95] | CAGTATCGGTTGGGAAGGGCGATCCGTTGTAA |
| 137 | 23[128] | 25[127] | CAGCTTTCAAAGCGCCATTCGCCAATGCCTGC |
| 138 | 23[160] | 24[144] | TGATTGTTGATGGCAATTCATCAATGCCGGA |
| 139 | 23[192] | 25[191] | AGGGTTAGCGGAATTATCATCATAGATAATAC |
| 140 | 23[224] | 25[223] | GCACGTAATTTTGCGGAACAAAGAACTTTACA |
| 141 | 24[79] | 22[80] | GCCTCTTCCTGCCAGTTTGAGGGTCTCCGT |
| 142 | 24[111] | 22[112] | GCGCAACTGCCTCAGGAAGATCGCATTAAATG |
| 143 | 24[143] | 23[159] | AACCAGGCCGGCACCGCTTCTGGTATAATCC |
| 144 | 24[175] | 22[176] | TATCAGATTGGATTATACTTCTGAATACCAAG |
| 145 | 24[207] | 22[208] | AGAAGGAGAACCTACCATATCAAAAACAAT |
| 146 | 24[239] | 22[240] | CATTATCAAACAGAAATAAAGAAAATATACAG |
| 147 | 25[64] | 27[63] | CGCCAGGGCATACGAGCCGGAAGCGAGCTAAC |
| 148 | 25[96] | 27[95] | AACGACGGGAAATTGTTATCCGCTCTGCCCGC |
| 149 | 25[128] | 27[127] | AGGTCGACTTCGTAATCATGGTCACCAGCTGC |
| 150 | 25[160] | 26[144] | ACTAATAGAAAATATCTTTAGGAGGGTACCG |
| 151 | 25[192] | 27[191] | ATTTGAGGCAACAGTTGAAAGGAAAAAACAGA |
| 152 | 25[224] | 27[223] | AACAATTCCCCTCAATCAATATCTCGCCTGCA |
| 153 | 26[79] | 24[80] | CCACACAATTTTCCCAGTCACGAGGTGCGG |
| 154 | 26[111] | 24[112] | TCCTGTGTCCAGTGCCAAGCTTGCTTCAGGCT |
| 155 | 26[143] | 25[159] | AGCTCGAATCTAGAGGATCCCCGCACTAACA |
| 156 | 26[175] | 24[176] | GGTTATCTATTAGAGCCGTCAATATTCCTGAT |
| 157 | 26[207] | 24[208] | TGGCAAATATTTAGAAGTATTAGAACCACC |
| 158 | 26[239] | 24[240] | ATATCAAAGACAACTCGTATTAAATTGAGTAA |
| 159 | 27[64] | 29[63] | TCACATTACACCGCCTGGCCCTGACCCCAGCA |
| 160 | 27[96] | 29[95] | TTTCCAGTCCAGTGAGACGGGCAAGTTCCGAA |
| 161 | 27[128] | 29[127] | ATTAATGACGTATTGGGCGCCAGGAAAGAATA |
| 162 | 27[160] | 28[144] | CCGAACGACCTAAAACATCGCCATGGGAGAG |
| 163 | 27[192] | 29[191] | GGTGAGGCGGCTATTAGTCTTTAATCAATCGT |
| 164 | 27[224] | 29[223] | ACAGTGCCAGAATACGTGGCACAGCAGATTCA |
| 165 | 28[79] | 26[80] | TTGCCCTTATTGCGTTGCGCTCACACAATT |
| 166 | 28[111] | 26[112] | TCTTTTCACGGGAAACCTGTCGTGTAGCTGTT |
| 167 | 28[143] | 27[159] | GCGGTTTGATCGGCCAACGCGCGTAAAAATA |
| 168 | 28[175] | 26[176] | CTGATAGCACCACCAGCAGAAGATTTGAGGAA |
| 169 | 28[207] | 26[208] | TTTTGAATGGTCAGTATTAACACGGTCAGT |
| 170 | 28[239] | 26[240] | AAAGCGTAACGCTGAGAGCCAGCAAACCTCAA |
| 171 | 29[64] | 31[63] | GGCGAAAAATGGCCCACTACGTGAGGTGCCGT |
| 172 | 29[96] | 31[95] | ATCGGCAAGTCAAAGGGCGAAAAAAGGGAGCC |
| 173 | 29[128] | 31[127] | GCCCGAGAAGAGTCCACTATTAAAAGCCGGCG |
| 174 | 29[160] | 30[144] | ATGGAAATGCCATTGCAACAGGAATTCCAGT |
| 175 | 29[192] | 31[191] | CTGAAATGTGCTGGTAATATCCAGGAATCCTG |
| 176 | 29[224] | 31[223] | CCAGTCACGCCTGAGTAGAAGAACGGCCACCG |
| 177 | 30[79] | 28[80] | TCAGGGCGTCCTGTTTGATGGTGCAGCTGA |
| 178 | 30[111] | 28[112] | ACTCCAACAATCCCTTATAAATCAGTGGTTTT |
| 179 | 30[143] | 29[159] | TTGGAACATAGGGTTGAGTGTTGAAACGCTC |
| 180 | 30[175] | 28[176] | TACCGCCAACCTACATTTTGACGCTGCGCGAA |
| 181 | 30[207] | 28[208] | ATCGGCCTGATTATTTACATTGGACAATAT |
| 182 | 30[239] | 28[240] | CATCACTTACGACCAGTAATAAAACTGACCTG |
| 183 | 31[64] | 30[80] | AAAGCACTAAATCGGAACCCTAACCGTCTA |
| 184 | 31[96] | 30[112] | CCCGATTTAGAGCTTGACGGGGAAGAACGTGG |
| 185 | 31[128] | 31[159] | AACGTGGCGAGAAAGGAAGGGAATTAAAGGG |
| 186 | 31[160] | 30[176] | ATTTTAGACAGGAACGGTACGCCAAACAATAT |
| 187 | 31[192] | 30[208] | AGAAGTGTTTTTATAATCAGTGATCAAACT |
| 188 | 31[224] | 30[240] | AGTAAAAGAGTCTGTCCATCACGCAGTAATAA |
| 189 | 12[143] | 11[159] | AGCGTCCATAGTAAAATGTTTAGTTTATCCC |
| 190 | 9[160] | 10[144] | TCTTACCGGAAACAATGAAATAGCACTAACG |
| 191 | 7[192] | 9[191] | AGACAAAAGCAACATATAAAAGAAAAGTAAGC |
| 192 | 19[192] | 21[191] | AGACTACCCCCTTAGAATCCTTGAGATGAAAC |

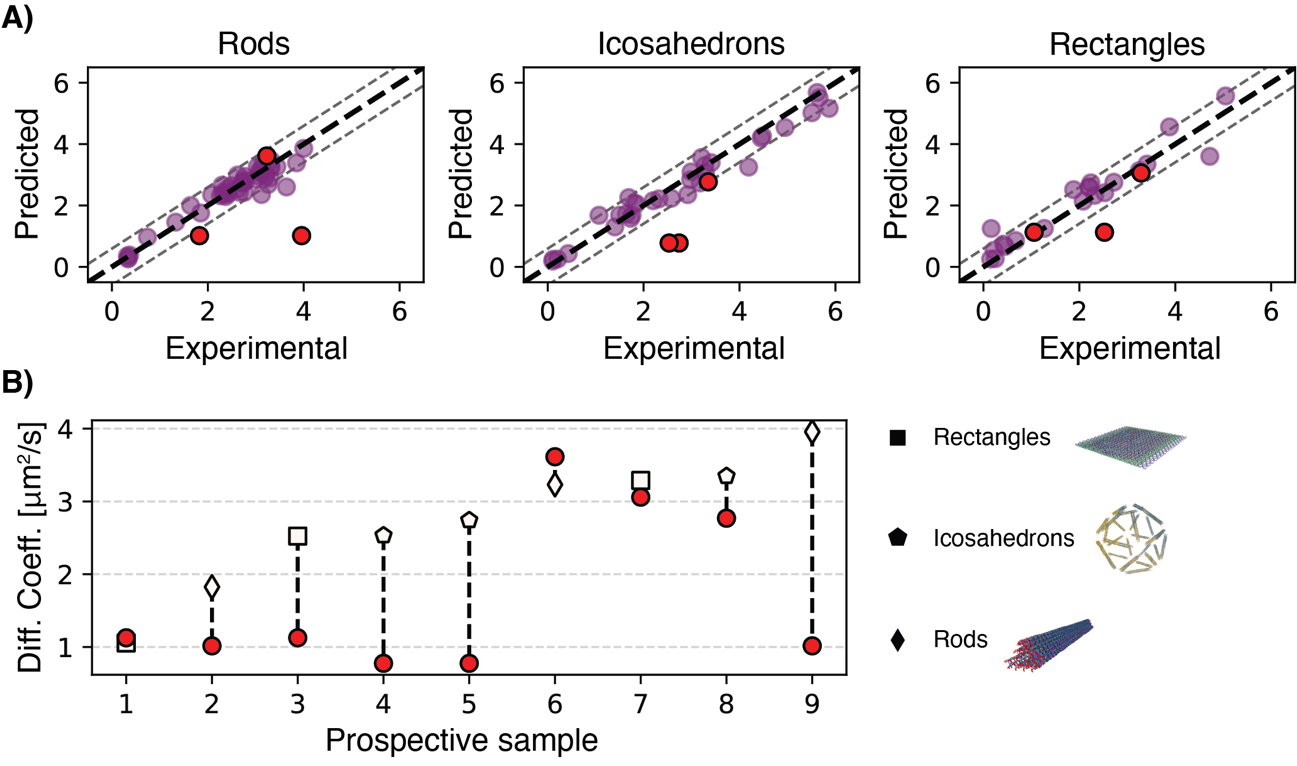

Figure S5. XGBoost analysis on individual shapes on the prospective dataset. A) Per-shape (i.e., Rods, Icosahedrons, and Rectangles) correlation plots showing in the background, without edge-colors in purple the prediction on the held-out set, and with red with black edge color in the foreground the prediction on the prospective dataset. The grey dotted lines delimit an interval of confidence that matches the average variability in experimental measurements. B) The tested prospective samples ranked based on the extent of their recorded diffusion coefficient (increasing from left to right). The dotted vertical line represents the distance between the predicted diffusion coefficient (circles) and the real experimentally recorded value (shaped data points).

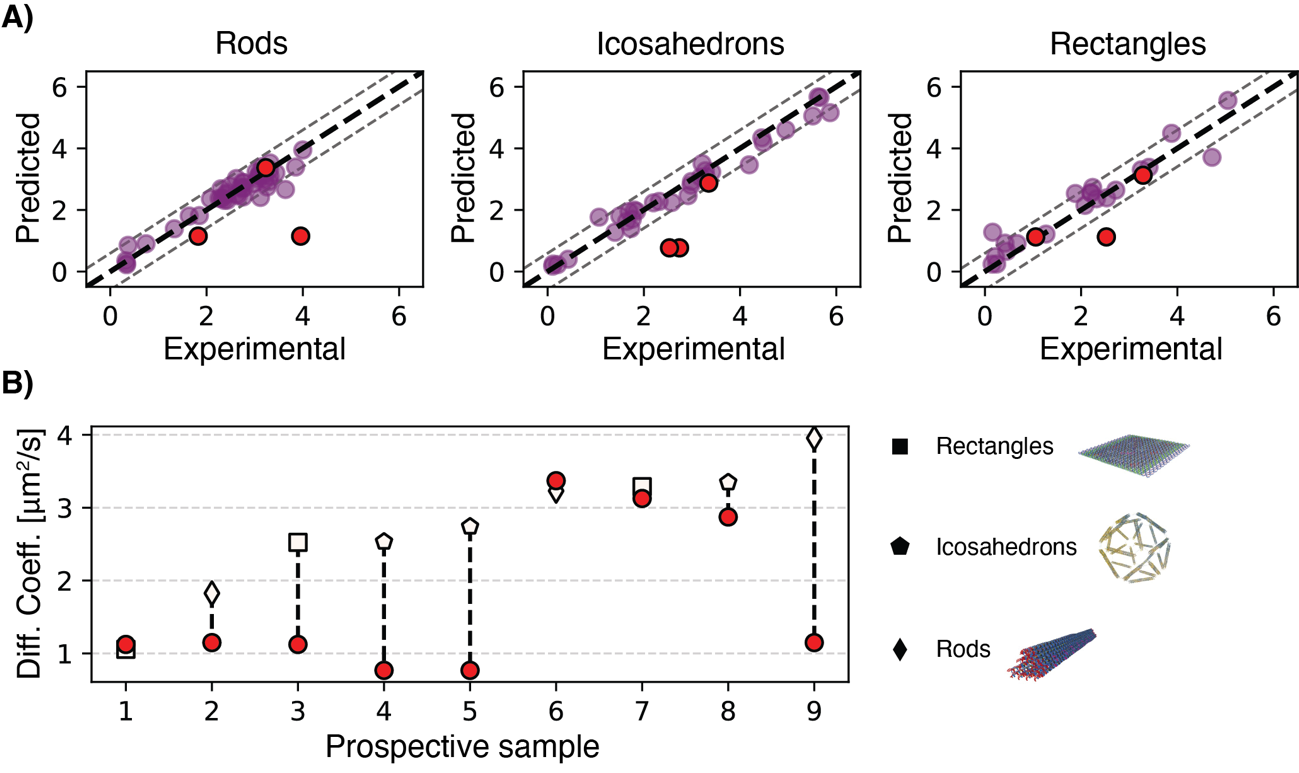

Figure S6. Random Forest analysis on individual shapes on the prospective dataset. A) Per-shape (i.e., Rods, Icosahedrons, and Rectangles) correlation plots showing in the background, without edge-colors in purple the prediction on the held-out set, and with red with black edge color in the foreground the prediction on the prospective dataset. The grey dotted lines delimit an interval of confidence that matches the average variability in experimental measurements. B) The tested prospective samples ranked based on the extent of their recorded diffusion coefficient (increasing from left to right). The dotted vertical line represents the distance between the predicted diffusion coefficient (circles) and the real experimentally recorded value (shaped data points).

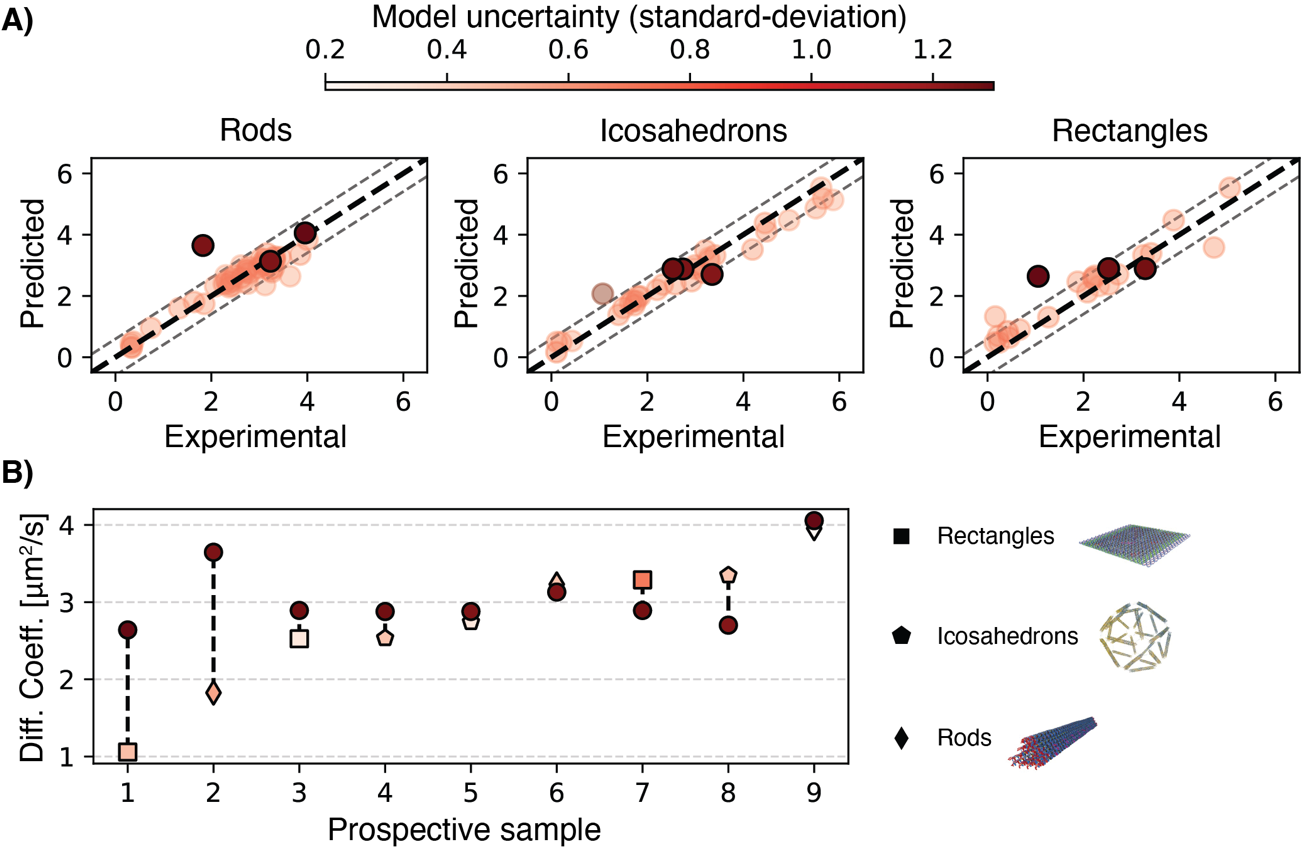

Figure S7. GPR-2k analysis on individual shapes on the prospective dataset. A) Per-shape (i.e., Rods, Icosahedrons, and Rectangles) correlation plots showing in the background, without edge-colors the prediction on the held-out set, and with black edge color in the foreground the prediction on the prospective dataset. The points red color intensity is proportional to the model predicted uncertainty. The grey dotted lines delimit an interval of confidence that matches the average variability in experimental measurements. B) The tested prospective samples ranked based on the extent of their recorded diffusion coefficient (increasing from left to right). The dotted vertical line represents the distance between the predicted diffusion coefficient (circles) and the real experimentally recorded value (shaped data points).
